# A molecular phylogeny of bedbugs elucidates the evolution of host associations and sex-reversal of reproductive trait diversification

**DOI:** 10.1101/367425

**Authors:** Steffen Roth, Ondřej Balvín, Osvaldo Di Iorio, Michael T. Siva-Jothy, Petr Benda, Omar Calva, Eduardo I. Faundez, Mary McFadzen, Margie P. Lehnert, Faisal Ali Anwarali Khan, Richard Naylor, Nikolay Simov, Edward H. Morrow, Endre Willassen, Klaus Reinhardt

## Abstract

All 100+ bedbug species (Cimicidae) are obligate blood-sucking parasites and well-known for their habit of traumatic insemination but the evolutionary trajectory of these characters is unknown. Our new, fossil-dated, molecular phylogeny estimates that ancestral Cimicidae evolved ca. 115MYA as hematophagous specialists on an unidentified host, 50MY before bats, switching to bats and birds thereafter. Humans were independently colonized three times and our phylogeny rejects the idea that the divergence of the two current urban pests (*Cimex lectularius* and *C. hemipterus*) 47MYA was associated with the divergence of *Homo sapiens* and *H. erectus* (1.6MYA). The female’s functional reproductive tract is unusually diverse and heterotopic, despite the unusual and strong morphological stasis of the male genitalia. This sex-reversal in genital co-variation is incompatible with current models of genital evolution. The evolutionary trait diversification in cimicids allowed us to uncover fascinating biology and link it to human pre-history and current activity.

## Introduction

Bedbugs (Cimicidae) are best known for the dramatic global re-surges of the pest, *Cimex lectularius* (the bedbug). The group actually comprises >100 described species (Usinger, 1966; Henry, 2009) of (secondarily) wingless blood-suckers that feed on mammals or birds and require transport by their hosts for dispersal. All members show traumatic insemination (Usinger, 1966). A phylogeny of bedbugs is required to solve four major unresolved biological puzzles. First, hematophagy hypothetically evolved in an ancestral opportunistic predator associated with vertebrate nests that took adventitious blood meals (Lehane, 2005; Weirauch et al., 2018). This scenario predicts that the ancestor should be a ‘host generalist’. However, all cimicid taxa currently considered basal (Usinger, 1966) are host specialists (Usinger, 1966, Ueshima, 1968). Moreover, bats are believed to be the ancestral host (Usinger, 1966), an assumption that requires testing because the oldest known cimicid fossil (100 MYA) (Engel, 2008) predates the oldest known bat fossil (Simmons et al., 2008) by ca. 50 MY.

Second, three cimicid species rely on humans as their main host: two have acquired urban pest status (Harlan et al., 2008). Specialized host use (one or few host species; specialists, S) in parasites is predicted to evolve by selection for resource efficiency in species with a broad host ‘portfolio’ (generalists, G) (Futuyma and Moreno, 1988; Poulin et al., 2006; Janz and Nylin, 2008; Hardy & Otto, 2014; Day et al., 2016; Hoberg and Brooks, 2008; Agosta et al., 2010; Park et al., 2018). However, in cimicids it is important to predict how, and how easily and rapidly, new species (such as humans) are accommodated in the host portfolio. Phylogenetic reconstructions of host-use assist in identifying factors that affect vertebrate host dynamics in human pre-history as well as in the context of current human activity including range alterations of wildlife by climate change (Pacifici et al., 2017), and by the livestock and pet trades.

A third question is whether the origin of human-associated parasites can be traced back to independent evolution on *Homo sapiens* and *H. erectus* that diverged 1.6 MYA, and the failure of the parasites’ gene pools to merge upon extinction of *H. erectus* ca. 100,000 years ago. This idea was suggested by Ashford for the common and the tropical bedbugs, *C. lectularius* and *C. hemipterus*, among four other parasite species pairs on humans (Ashford, 2000). However, the only empirical evidence (from the head louse) is conflicting (Reed et al., 2004; Kittler et al., 2003). The fact that *C. lectularius* and *C. hemipterus* are able to mate with each other (albeit without producing offspring) (Coetzee et al., 1995) suggests a recent divergence of the two taxa. By contrast, accommodating all the speciation events that happened within the *C. lectularius* and *C. hemipterus* clades after they diverged (Balvin et al., 2015) would require unusually high speciation rates.

Fourth, the traumatic insemination of bedbugs, the obligatory copulatory wounding of females by males (Usinger, 1966; Reinhardt et al., 2003, 2014; Tatarnic et al., 2014; Stutt and Siva-Jothy, 2001; Morrow and Arnqvist, 2003), is a commonly cited example of the evolutionary conflict between males and females. The fitness costs arising from traumatic insemination selected for a unique female ‘defense’ organ, the spermalege (Morrow and Arnqvist, 2003; Reinhardt et al., 2003; Stutt and Siva-Jothy, 2001). Unlike the female genitalia proper, the spermalege does not function in egg-laying and so is free to evolve in response to variation in male mating traits. This organ is highly variable across species, showing species-specific location on the body (heterotopy) and considerable variation in anatomical complexity (Usinger, 1966). The variation in cimicid spermalege structure or position has not been compared with models of genitalia evolution (Eberhard, 1985; Hosken and Stockley, 2004; Brennan and Prum, 2014) due to the lack of a phylogeny. Understanding the evolutionary trajectory of the spermalege would also impact on classical systematics because the lack of a spermalege in *Primicimex* has been assumed to represent the evolutionarily ancestral state (Usinger, 1966; Schuh and Slater, 1995). While possible, it is not consistent with the idea that *Bucimex*, which has a spermalege, is *Primicimex*’s sister group (Usinger, 1966; Schuh and Slater, 1995). A molecular phylogeny can test this ‘sister group’ hypothesis and indicate whether the taxon is indeed basal within the Cimicidae.

Fresh cimicid material, collected over 15 years (Supplement 1), has allowed us to reconstruct and date the first molecular phylogeny of the group, providing insights into the evolution of bedbugs and hematophagy, patterns of host utilization and a unique analysis and understanding of the evolution of female genital variation.

## Results and Discussion

### The molecular phylogeny

The consensus tree (Fig. 1) i) shows the Cimicidae are monophyletic and firmly placed within the Cimicomorpha (Weirauch et al., 2018; Schuh et al, 2009; Jung and Lee, 2012), ii) provides robust resolutions of other debated relationships (Fig. 1), including the paraphyly of the martin bugs (Balvin et al., 2015; Jung and Lee, 2012) and iii) exposes the geographic structure expected for wingless, poor dispersers (Fig. 1), even though most colonization events are not recent. Our consensus tree iv) robustly identified *Primicimex*+*Bucimex* as a monophylum (supporting morphological arguments – Usinger, 1966) and as the sister of the remaining extant Cimicidae (basal lineage), solving a long-standing problem in insect systematics (Schuh and Slater, 1995; Schuh et al., 2009; Jung and Lee 2012). It also shows that the lack of a spermalege (and of a mycetoma – Usinger, 1966) in *Primicimex* cannot readily be interpreted as an ancestral state because *Bucimex* possesses both organs. Instead, our phylogeny suggests two new hypotheses for the evolution of the spermalege. *Primicimex* shows dorsal insemination but its extant sister genus *Bucimex* shares the ventral site of insemination with the sister clades (Usinger, 1966; see below). Therefore, either i) a change in *Primicimex* from ventral to dorsal intromission site was paralleled by a loss of the spermalege in *Primicimex*, or ii) the ventral position of the spermalege evolved independently in *Bucimex* and in the rest of the Cimicidae. Given that two other derived species not studied here (*Rusingeria transvaalensis, Crassicimex pilosus*) independently lack a spermalege (Usinger, 1966), and that the transition from ventral to dorsal insemination is rare (see below), the secondary loss of the spermalege in *Primicimex* is the most likely scenario.

**Fig. 1.**
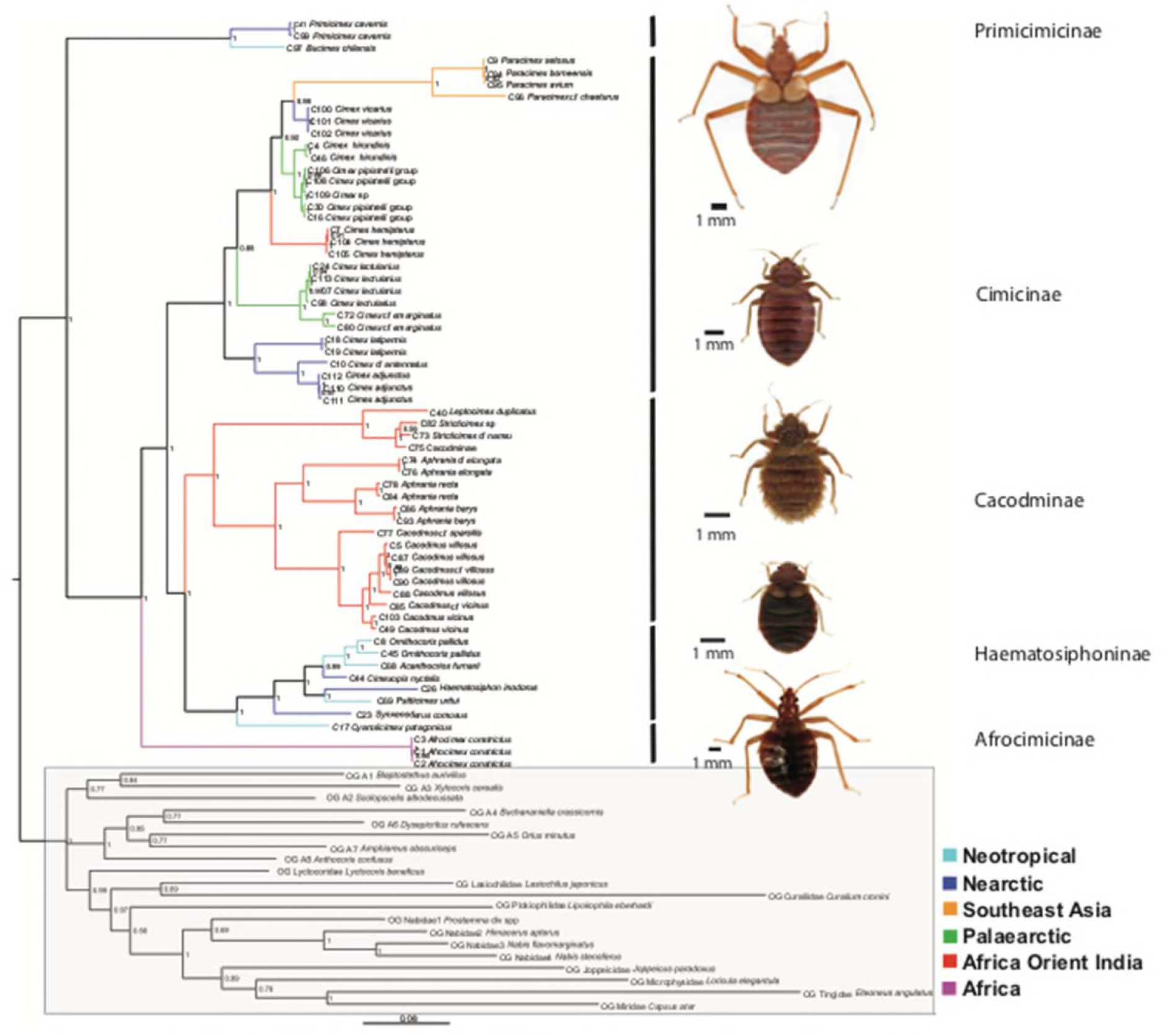
Phylogeny of the bedbug family (Cimicidae). Bayesian consensus tree based on four genes showing the biogeographical distribution and classical taxonomy at the subfamily level. Photographs show typical representatives of each subfamily. Numbers beside the nodes indicate posterior probability values. The tree reveals the monophyly of several debated taxa such as i) *Primicimex* + *Bucimex*, ii) the Nearctic/Neotropical Haematosiphoninae, iii) *Paracimex* + *Cimex*(and confirms earlier findings that *Cimex* is paraphyletic incorporating the former genus *Oeciacus* (Balvin et al., 2015). The branch lengths scale represent the number of estimated nucleotide substitutions per site. Sample codes refer to Supplement 1. Sequences of outgroups, boxed in shaded grey, were taken from GenBank.

The underlying data (**Additional Data 1**) are available at ***dryad*** (see additional files; currently uploaded for review).

### Enigmatic ancestral host and multiple colonization events of bats

Independently dating the phylogenetic tree using a fossil from the related family Vetanthocoridae (152 MYA) (Yao et al., 2007) rejects the widely-held view (Usinger, 1966) that the Cimicidae evolved on bats. Our mean estimate of 115 MYA (74–170, 95 % highest posterior density (HPD) interval) for the stem of the Cimicidae supports the idea of a minimum age of the group of 100 MYA based on fossil evidence (Engel, 2008). The origin of the Cimicidae crown group with a mean of 93.8 (56–137 95% HPD) MYA is placed 30–50 MY before bats are known to have evolved (Simmons et al., 2008; Teeling, 2005; Agnarsson et al., 2011; Lei and Dong, 2016) (Fig.2). This estimate appears robust: Employing the oldest known cimicid fossil as an additional calibration point shows the stem species evolved about 122 MYA (111–150 MYA, 95 % HPD, relaxed molecular clock estimation of lineage divergence points within the family) and the crown diverged from about 102 MYA (91–114 MYA, 95 % HPD) (Fig. 2).

**Fig. 2.**
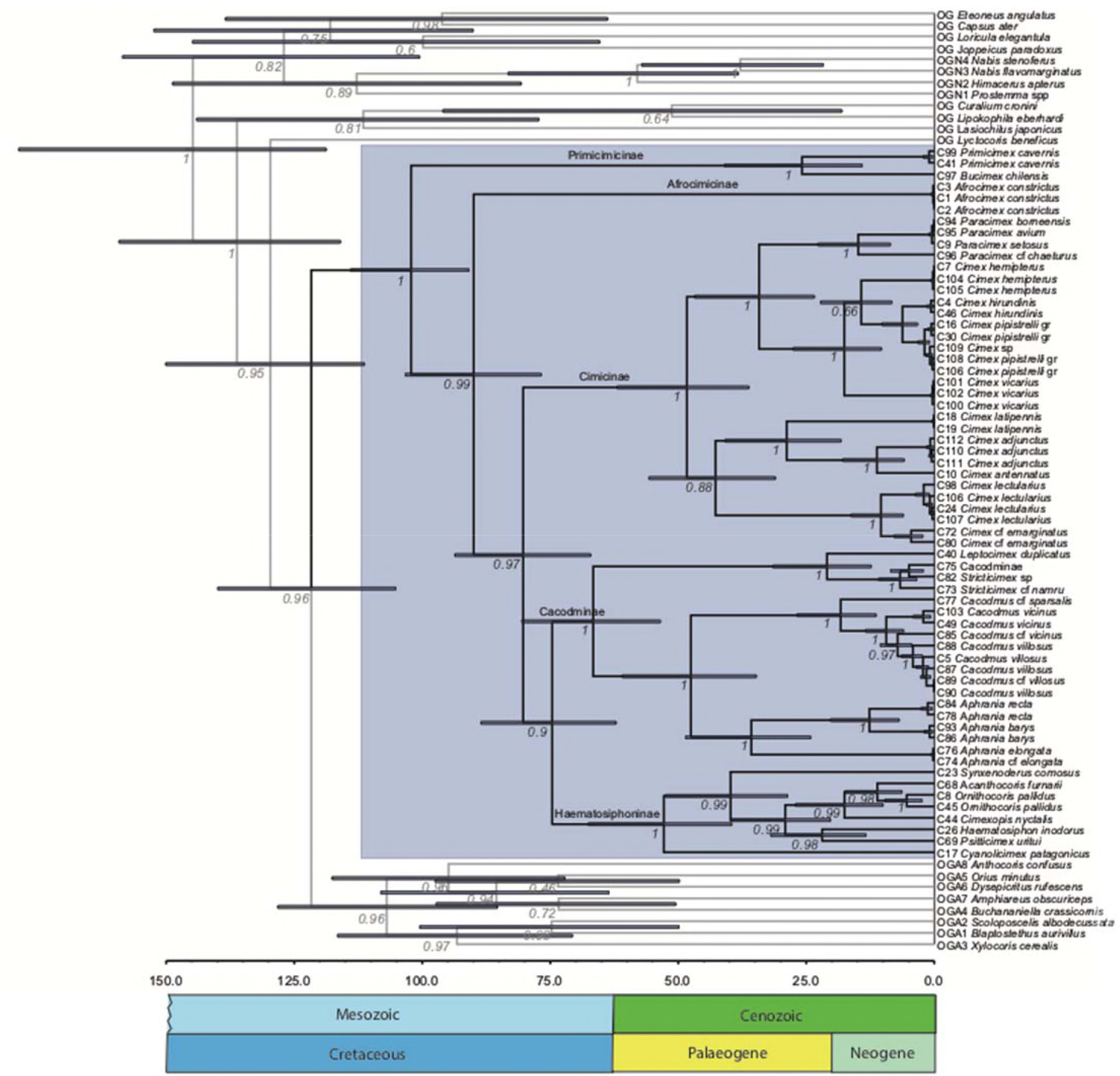
Chronogram of the bedbug family (Cimicidae). Bayesian consensus tree of the Cimicidae and selected outgroup taxa in relation to geological age (MYA) (x-axis). A relaxed clock model (Drummond et al., 2012) was used to date the tree based on two calibration points, fossil Vetanthocoridae (152 MYA) (Yao et al., 2006) and the oldest known fossil cimicid (100 MYA) (Engel, 2008). Numbers below nodes represent Bayesian posterior probability values, blue bars represent 95% highest posterior density intervals of the time estimates in million years (MYA). Scale in millions of years. The Cimicidae are boxed in shaded blue. ‘gr.’ stands for group, a taxonomic aggregate.

The underlying data (**Additional Data 1**) are available at ***dryad*** (see additional files; currently uploaded for review).

All four ancient bedbug lineages predate the evolution of bats (Fig. 2) but were reconstructed to ancestrally used bat hosts. This suggests that bats were colonized several times independently (Fig. 3A), unless the evolutionary origin of bats (Simmons et al., 2008; Teeling, 2005; Agnarsson et al., 2011; Lei and Dong, 2016) has been grossly underestimated.

**Fig. 3.**
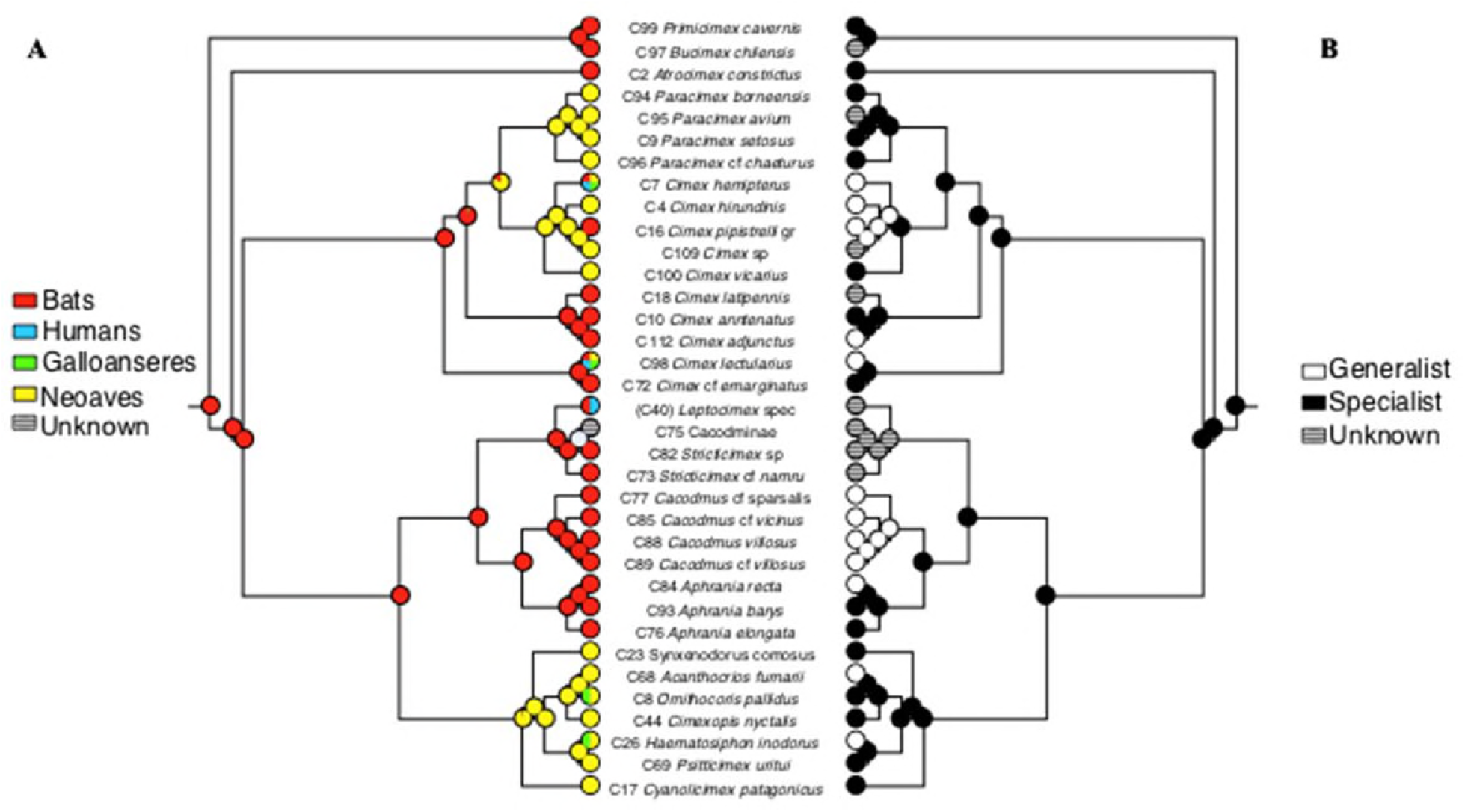
Ancestral bedbug hosts. Mirror trees showing (A) Systematic host groups, and (B) As host specialist or generalist. The Bayesian consensus tree was used with the trace character state function in the software Mesquite (Maddison and Maddison, 2017). Proportional probabilities of ancestral hosts were reconstructed using likelihood and a one parameter Markov model with state changes rates estimated from the data (Lewis, 2001). In A), colors indicate different host types reported (Usinger, 1966; Ueshima, 1968; Di Iorio et al., 2013); Országh et al., 1990). In B) specialized (black) or generalist (white) host use were reconstructed with (unordered) parsimony. Separate analyses with BayesTraits confirmed specialized host use as the ancestral state for all Cimicidae. The result did not change if the two lineages with the highest uncertainty about their ancestral state, i.e. Cimicinae and (Cacodminae + Haematosiphoninae) were analysed separately by setting all other clades to an unknown state of G or S (probabilities of bats as ancestral host 98%, and ancestral specialist 85% for the Cimicinae, and 96%, and 98%, respectively for the Cacodminae+Haematosophinae. *Leptocimex duplicatus* was analysed as *Leptocimex* spec. to demonstrate human host use in this genera. Results were identical if ancestral analysis host and specialization employed bats or bats +human).

The underlying data (**Additional Data 1**) are available at ***dryad*** (see additional files; currently uploaded for review).

To summarise, our independently dated molecular phylogenetic tree estimated the stem species of bedbugs at 115–122 MYA, well before the K-T mass extinction boundary, a key event in vertebrate diversification. The identity of the ancestral hosts remains a mystery but bats were colonized repeatedly.

**Evolution of hematophagy**. We clearly reject ancestral host generalism (G) in cimicids (Fig 3B), and therefore the evolution of hematophagy from facultative blood-feeding by ancestral predators (Lehane, 2005; Weirauch et al., 2018). This holds true if G were broadly defined by the phylogenetic distance of their hosts (Park et al., 2018) as using more than one of the four major, phylogenetically deeply diverged host groups of waterfowl (Galloanseres) and other birds (Neoaves), as well as bats (Chiroptera) and humans (Fig. 3A). It also holds true for a tighter definition of G that includes variability within taxonomic group (Park et al., 2018) as being those parasites recorded from more than three genera (Supplement 2). Therefore, hematophagy likely evolved before the Cimicidae, within the true bugs (Heteroptera), in insects that were already specialists. This result is consistent with the view that the specialist blood-sucking Polyctenidae are the sister group of the Cimicidae (Schuh and Slater, 1995).

### Pattern of host shifts

Defining species along the host specialist (S)/host generalist (G) axis depends on the specialization metrics and on recording intensity (Futuyma and Moreno, 1988; Poulin et al., 2006; Janz and Nylin, 2008; Hardy & Otto, 2014; Day et al., 2016; Hoberg and Brooks, 2008; Agosta et al., 2010; Park et al., 2018). Of the 29 species on our tree that allow a classification, most (24/29, 83%) are S (broadly defined – figs 3A), or 55% (15/27), when more tightly defined (Supplement 2). Five cimicid species on our molecular tree are G (broadly defined). Of the species not represented on our tree, five more were classifiable, three G and two S (Usinger, 1966).

Host shifts by the ancient bat specialists were common since most extant bat-parasitic cimicid lineages evolved before their extant hosts lineages (Supplement 3). Host switches also occurred at least three times independently from bats to birds (Fig. 3A). Our host reconstruction indicates (Hoberg and Brooks, 2008; Agosta et al., 2010; but see Hafner et al., 1994) that parasite diversification is not generally driven by co-speciation with either bat or bird hosts (Supplements 4 and 5). Together these observations suggest that the extant pattern of G/S distribution in cimicids is the result of evolutionarily dynamic host transitions.

When examining host transitions at all 31 subterminal nodes on our tree that are classifiable as G or S, we found the highest number (9/31, or 29%) involved host specialists switching host but staying specialist (S→S). Two nodes were G→S transitions (6%) and five (16%) were S→G transitions (or 7/31 (23%) if specialists are defined more strictly) (Figs. 3B, Supplement 2).

The paucity of G→S transitions, departing from the general pattern in mammalian parasites (Park et al., 2018), indicates that the “resource efficiency” hypothesis where host specialists (S) evolve from generalists (G) by fitness advantages on specific hosts (Futuyma and Moreno, 1988; Poulin et al., 2006; Janz and Nylin, 2008) rarely applied to cimicids. A modification of this idea, the “oscillation” hypothesis, proposes that maintained genetic variation or phenotypic plasticity allows S species to add hosts to their portfolio to become G again, depending on ecological opportunities (Hardy and Otto, 2014; Hoberg and Brooks, 2008; Park et al., 2018). This hypothesis is difficult to test as it allows for any number of S/G transitions. However, if S→G transitions were regularly oscillating, they should be evenly distributed across evolutionary time. By contrast, all seven S→G transitions were recent, between 10 and 20 MYA (cf. figs. 2,3B).

In case ancient host ranges are hard to reconstruct (Janz and Nylin, 2008), stochastic acceptance of unusual hosts, such as laboratory-forced host feeding might serve as an indicator of plasticity or genetic variation (Hoberg and Brooks, 2008). Such stochastic host use has so far only been recorded in G (Figs. 3, Supplement 2) but not in S species (Ueshima, 1968; K. Reinhardt, R. Naylor, M.T. Siva-Jothy, unpubl. data on *Afrocimex constrictus*) as required by the “oscillation hypothesis”. Outside the laboratory, they have occurred mainly during ecological opportunities created by humans, such as guano-mining, chicken-breeding or pet-keeping. Even if systematic laboratory screens of S reveals unusual hosts, there is little current evidence that host specialization in the Cimicidae is driven by selection for resource efficiency. The pattern is not consistent with the oscillation hypothesis either.

S→S transitions (host switches without extensions in host breadth, or “musical chairs” pattern –Hardy and Otto, 2014) are the common pattern in cimicids. Like S→G, S→S transitions might also be based on the ecological opportunities new hosts represent (Hoberg and Brooks, 2008; Agosta et al., 2010), such as after major (e.g. inter-continental) dispersal events (Park et al., 2018). For example, two of the three bat-to-bird host shifts concerned the Haematosiphoninae and *Paracimex* where bird hosts replaced bats, rather than having been added (Fig. 3) (the third situation cannot be reconstructed) and both examples also involved the colonization of another continent (South America and Southeast Asia). However, other S→S transitions are not related to inter-continental changes.

To summarise, several blood-feeding bedbug lineages specialized on bats in ancient times. Subsequent host shifts in the Cimicidae were frequent and the switches between hosts as well as expansions of the host portfolio that can be explained, are related to the ecological opportunities that human activity or inter-continental dispersal provided.

### Human colonization and Ashford’s hypothesis

Three bedbug species were reported to routinely use humans as hosts (*C. lectularius, C. hemipterus* and *Leptocimex boueti*) (Usinger, 1966, Lehane, 2005; Harlan et al., 2008). All are G, all are recent and all represent expansions of the host portfolio, not replacements, i.e. they represent the somewhat more unusual S→G transitions among mammalian parasites (Park et al., 2018) (Fig. 3). All colonization of humans is non-randomly captured by these S→G transitions, which represent just 16% (or broad definition: 23%) of all transitions [Fisher’s exact test, *P*=0.0022 (or broad definition of G: *P*=0.0078)]. Thus, humans represent an important, nonrandom, target for specialist cimicid species to expand their host portfolio.

Our phylogenetic tree reveals that all three evolutionary events of human host use occurred independently (Fig. 3A). This notion, in concert with the finding that the *C. hemipterus*and *C. lectularius* lineages diverged ∼47 MYA, clearly rejects Ashford’s (2000) hypothesis, which predicts a divergence around 1.6 MYA to coincide with the split between the *H. sapiens* and the *H. erectus* clades. Since we identify *C. lectularius* as belonging to a bat-associated lineage and *C. hemipterus* to a bird-parasitic lineage (Balvin et al., 2015; Fig 3A), Ashford’s idea would additionally require a series of independent host shift from *Homo* linages to birds and bats. With one species pair of human parasites showing ambivalent support for this hypothesis (lice: Kittler et al., 2003; Reed et al., 2004) and one not (cimicids), Ashford’s idea should be re-tested by dating the split of other species pairs of human parasites.

*C. lectularius* has been hypothesized to have colonized humans, or *H. sapiens*, when ancient man started to become a cave-dweller (Usinger, 1966). However, our analysis shows all clades parasitizing humans had diverged at least 5–10 MY before the oldest known *Homo* species (White et al., 2009; Spoor, 2015). The coexistence of several lineages of hominids in space and time (Spoor, 2015) allows for several transmission scenarios and host shifts. However, because bedbugs are not known from other extant hominids, or indeed other primates, colonization took place primarily in the hominin lineages. Thus, no matter when hominids first entered caves, bat-and bird-parasitizing *C. lectularius* were already there, ready to exploit incoming opportunities. Thus, the fact that bat and human-associated lineages of *C. lectularius* diverged between 99,000–867,000 years ago (Balvin et al., 2012) provides a hint of when humans acquired *C. lectularius*, but not which of the *Homo* lineages, and neither whether cave-dwelling was the initial driver for contact. Similar questions should be asked for *C. hemipterus* and *L. boueti* but it is clear the transmissions track suggested by Ashford is too simplistic.

### Diversification of the female copulatory organ

The female copulatory organ, the spermalege, functions to offset costs of traumatic insemination (Reinhardt et al., 2014; Tatarnic et al., 2014; Stutt and Siva-Jothy 2001; Michels et al., 2015; Benoit et al., 2011). Unlike genitalia, the spermalege is not evolutionarily constrained by egg-laying and so can be expected to show even more rapid co-evolutionary responses to copulation. Consistent with this idea, the spermalege ranges in anatomical complexity from being absent, to a simple cuticular invagination, to being duplicated, or even manifest as a secondary, epithelium-lined paragenital tract. However, unlike the striking parallel found in a related group of plant bugs (Tatarnic and Cassis, 2010), the spermalege of cimicids shows a degree of heterotopy (West-Eberhard, 2003) across species that is unmatched in the animal kingdom (Supplement 6). The organ can occur dorsally or ventrally, left, right or in the middle, and on, or across, one or several of various segments along the body axis (Usinger, 1966) (Figs 4, Supplements 7 and 8). A change in spermalege position (Figs 4, Supplement 7) is associated with a change in biogeographic (continental) distribution in only 7/16 cases (Supplement 8). Neutrally accumulated variation in allopatry is therefore unlikely to explain this aspect of female ‘genital’ variation.

**Fig. 4.**
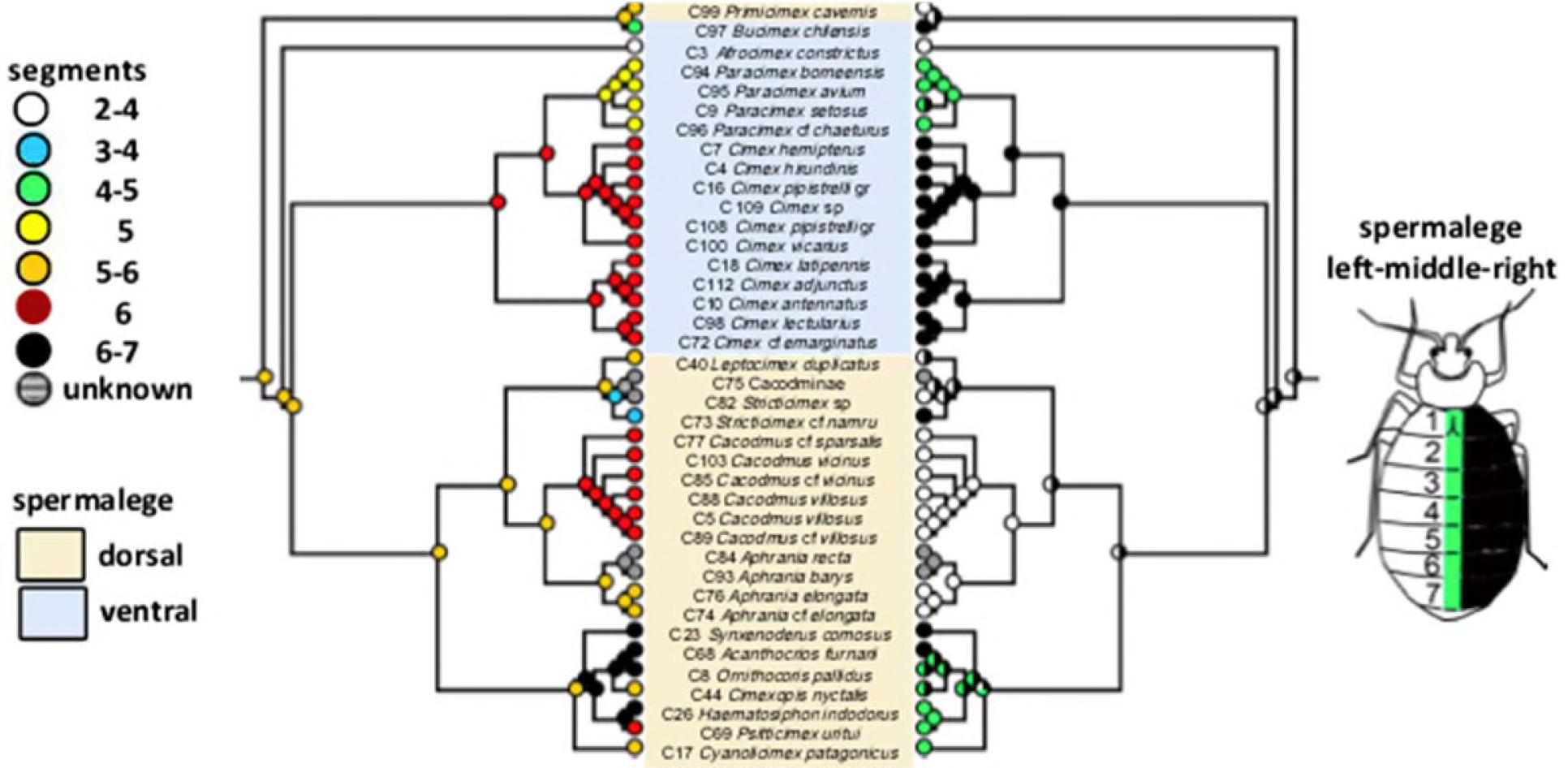
Heterotopy of the spermalege, the female copulatory organ in bedbugs. The spermalege is a defense organ against traumatic insemination and situated in a species-specific manner on the dorsal or ventral surface of the abdomen (central panel), on the left, right or central (mid) position of the body (right panel) and positioned along the cephalo-caudal axis (left panel) (Usinger, 1966). *Primicimex* lacks a spermalege but the corresponding segments of intromission are indicated. The ancestral states of the spermalege were reconstructed based on a Bayesian consensus tree using Mesquite parsimony, with probabilities generated from BayesTraits.

The underlying data (**Additional Data 1**) are available at ***dryad*** (see additional files; currently uploaded for review).

Several adaptive models explain the diversification of female genitalia. First, under classical sexual selection males diversify and female genitalia morphologically ‘follow’ the cross-species variation in male genitalia (Eberhard, 1985; Hosken and Stockley, 2004; Brennan and Prum, 2014). However, the lack of congruence between the small morphological variation in size and curvature of the left-asymmetric intromittent organs in males (Usinger, 1966; Eberhard, 2004; Supplement 7) and the huge diversity in complexity (Usinger, 1966) and heterotopy (Supplement 7) is not consistent with this idea.

Second, a mismatch of male and female genitalia can lead to traumatic insemination (Reinhardt et al., 2014), cause sexual conflict (Stutt and Siva-Jothy, 2001) and fuel coevolutionary cycles (Lessells, 2006; Gavrilets, 2014; Crespi and Nosil, 2013; Arnqvist and Rowe, 2002) between male and female copulatory organs. This pattern is observed in the traumatically inseminating plant bugs (Tatarnic and Cassis, 2010). However, the striking morphological stasis of male genitalia observed in cimicids is not consistent with males perpetually imposing sexually antagonistic evolution on females that result in fluctuating cycles of escalation and de-escalation at the morphological level (Arnqvist and Rowe, 2002). In addition, female heterotopy is ancient, not constantly (perpetually) evolving. On average, a positional change (heterotopy) occurred at 39.25 ± 26.8 MY (N=16), whereas nodes without heterotopy occurred at 17.15 ± 24.1 MY; N=20 (t = 2.571, df = 30.63, P = 0.0152), despite including the oldest split (Fig. 4). Of the 24 penultimate nodes (where most changes may be expected if coevolution is perpetual), only 9 involve any change in spermalege position.

Genital traits may also diversify morphologically if females tolerate, rather than resist, sexual conflict (Michels et al., 2015). However, again, this idea requires similar male and female variation at least some of the time (Michels et al., 2015) and therefore, does not apply here. The third model, called Buridan’s ass (Gavrilets and Waxman, 2002) suggests that female polymorphism evolves in response to sexual conflict because female mating damage is lower in divergent female morphs than in the original morph. Males, then condemned to low mating success between the diverged female mating morphs, experience reduced diversifying selection on their reproductive traits (‘jack of all trades, master of none’) (Gavrilets and Waxman, 2002). Whilst the pattern in cimicids is broadly consistent with this, a more detailed examination reveals some issues. Buridan’s ass requires disruptive selection, i.e. the novel female mating morphs do not include the ancestral state (otherwise the female morph cannot provide a female benefit and the male is not ‘caught in the middle’) (Gavrilets and Waxman, 2002). Hence, this model predicts that sister taxa should always be at least two character states apart from one another in heterotopy. Six out of 19 changes were of such a nature, despite three possible ‘axes’ of change but eight changes concerned a ‘next’-state character change that cannot occur under Buridan’s ass (Supplement 8). We conclude that ancient spermalege variation is maintained but not all changes were driven by the Buridan’s ass model.

One possibility left to explain the diversity in the site of the female paragenitalia in cimicids is to combine two observations: i) copulatory trauma also arises when male copulatory behavior and female morphology ‘mismatch’ (Reinhardt et al., 2014) and ii) male mating position frequently changes during evolution, generally driving female morphology in insects and has generated some ancient variation (Huber et al., 2007). Therefore, it may be male mating behavior that is under classical diversifying sexual selection in cimicids, even if male genitalia are under stabilizing selection. The change in female spermalege position may then respond in order to mitigate the cost of traumatic insemination imposed by changes in male mating position. This idea is consistent with other phenomena in the Cimicomorpha (supplementary discussion).

### Conclusion

In summary, our phylogenetic reconstruction i) shows that bedbugs evolved before their assumed primary bat hosts and colonized them on several occasions subsequently, ii) supports the view that generalism can evolve when ecological opportunities arise even after long periods of specialization, iii) shows that all colonization of human hosts were such cases. Our phylogenetic tree also iv) rejects the Ashford hypothesis, and v) suggests that male mating behavior may have driven female genitalia heterotopy without significantly affecting male genital evolution.

## Material and methods

### Material and sample origins

We obtained material from three main sources. First, we contacted the major natural history museums in the world as several species are known only from a single collection from their type locality. However, most of this material dated from the 1960s and 1980s that did not allow DNA analysis. In fact, museum material from only two species was useful for the molecular analysis, one of which was subsequently obtained otherwise. Second, between 2002 and 2015, we contacted researchers with requests for specimens. Researchers working on cimicids provided material from 10 species. We also contacted approximately 500 researchers that work in caves, on cave-dwelling bats or other putative bedbug hosts, such as swallows and swiftlets. Approximately half the people responded, and about 30 respondents promised to send material. From those who did send material, an extra 18 species were obtained. Third, between 2000 and 2014, the authors undertook field trips to obtain material, adding 10 species. This resulted in a total of 38 species, of which 34 species from 62 localities yielded sufficient DNA for the analysis (Supplement 1). Unfortunately, existing *Latrocimex* material from Brazil (Graciolli et al., 1999) was not at our disposal to be analyzed.

### Taxon sampling

In total, 34 species of Cimicidae were analysed, representing 17 genera from 5 out of 6 currently recognized subfamilies (Usinger, 1966). The most closely related families were chosen as outgroups: Nabidae, Anthocoridae, Plokiophilidae, Microphysidae, Curaliidae, and Joppeicidae (Weirauch et al., 2018; Schuh and Slater, 1995; Schuh et al., 2009; Jung and Lee 2012), except the Polyctenidae (for which we obtained no material). We also included representatives of two more distant outgroups, the Tingidae and Miridae. All outgroup taxa sequences were obtained from GenBank (Supplement 1).

### DNA extraction, PCR amplification, and DNA sequencing

Nuclear and mitochondrial genomic DNA was extracted from 70–96% ethanol-preserved specimens using a QIAGEN DNAEasy blood and tissue kit (Qiagen Inc., Hilden, Germany) following the manufacturer’s instructions and standard methods for DNA extraction and purification. If high-quality amplicons were not acquired, a set of ambiguous primers with universal sequencing adaptors was used (table S3). The total volumes of PCR reactions were 10 μl (0.25 μl Promega GoTaq Flexi DNA Polymerase (5 U/μl); ddH2O; 5x Colorless buffer; 2 mM MgCl2; 0.2 mMdNTP; 0.5 μM of each primer), with 1–2 μ DNA template. PCR thermal conditions are shown in Supplement 1. PCR products were purified using ExoSAP-IT. Sequencing reactions for both strands of the amplified genes were performed using BigDye Terminator v3.1 Cycle Sequencing Kit (Applied Biosystems). Products were sequenced using Applied Biosystems automated sequencer. Sequence contigs were assembled in Sequencher v. 4.5 (Gene Codes, Ann Arbor, Michigan).

Five nuclear and mitochondrial molecular markers were amplified comprising fragments of four gene regions (COI, 16S rDNA, 18S rDNA, 28S D3 rDNA) and selection models were selected in MEGA v.6 based on Akaike Information Criterion (Supplement 13).

### Sequence alignments

Alignment was conducted using the MUSCLE (Edgar, 2004) algorithm implemented in MEGA v. 6 (Tamura et al., 2013) with the following settings: –400 gap opening penalty, –50 gap extension penalty. We used GBlocks V.0.91b (Castresana, 2000) to test and where required to eliminate poorly aligned positions in the original alignments and used this dataset for an alternative analysis (Supplement 14).

Since saturation in substitutions can lead to incorrect phylogenetic inferences (Swofford, 1996), the positions 1–3 were evaluated for substitution saturation by DAMBE V 5.2.13 (Xia and Xie, 2001) in the whole dataset. Saturation was not observed for any but the third position in only the COI dataset. As there was no conflict in topology of the separate gene trees (see below) we run the analysis with all three positions. We tested several available specimens from *C. lectularius* for mitochondrial heteroplasmy (Robinson et al., 2015), but detected none (N=8).

### Phylogenetic analyses

Five molecular markers were amplified comprising fragments of four gene regions (COI, 16S rDNA, 18S rDNA, 28S D3 rDNA). Since there was no sequence overlap of the two 18S rDNA fragments in some taxa, the two fragments were treated as separate markers (called 18S part1 and part2) in all analyses. Models of evolution for each marker were selected in MEGA v.6 based on Akaike Information Criterion (Supplement 13). Preliminary analysis of single gene sets was unable to recover stable clades at different depths of the tree but did not show any conflict among the separate gene trees (Supplement 10). Therefore, phylogenetic Bayesian analyses (BA) were conducted on the concatenated data set in MrBayes 3.2. (Drummond et al., 2012). Model parameter values for the partitions were estimated independently using the “unlink” command and relative site-specific rates for all gene fragments were estimated by setting the prior for “ratepr” to “variable”. For all analyses, Markov Chain Monte Carlo (MCMC) sampling was conducted with two independent and simultaneous runs for 10,000,000 generations. Trees were saved every 1000 generations. Likelihood values and effective sample size were observed with Tracer v1.4 (Rambaut et al., 2014), and all trees sampled before the likelihood values stabilized were discarded as burn-in. Stationarity was reassessed using the convergence diagnostics in MrBayes (i.e., the average standard deviation of split frequencies [values <0.01] and the potential scale reduction factor [values ≈1.00]). A burn-in of 25% of all sampled trees was sufficient to ensure that suboptimal trees were excluded. The remaining trees were used to construct a 50% majority rule consensus tree.

Bayesian and other trees were formatted for presentation using either TreeView (Win32) 1.6.6 (Page, 1996), FigTree 1.4.1 (Rambaut, 2014), or Mesquite 3.2. (Maddison and Maddison, 2017) In order to test the robustness of our dataset we performed additional analyses using different outgroups. We found no effect on topology and support values for the ingroup clades (results not shown except Supplement 11 for a selection of the closest outgroup taxa). *Paracimex* showed no long branch attraction (Balvin et al., 2015).

In order to compare the tree from Bayesian inference with Maximum Likelihood (ML) analysis we ran the same partitioned dataset by using RAxML 7.4.2. (Stamatakis, 2006). Since RaxML does not allow the use of mixed nucleotide models, we used the GTRGAMMAI for all partitions. ML with rapid bootstrap was performed in 1000 iterations and obtained bootstrap values were placed on a consensus tree (Supplement 13).

### Molecular Dating

We used Beast 1.8.4 (Drummond et al., 2012) with 70 sequences, including eight outgroups to infer the divergence dates of the sequences under a Yule speciation process (a pure birth process) and an uncorrelated relaxed molecular clock (Drummond et al., 2012).

First, we constrained the Cimicoidea as a monophyletic group and used a lognormal prior mean age of 152.2 million years (MY) with standard deviation 0.2 MY as calibration point for the group based on a fossil flower bug (Heteroptera: Cimicomorpha: Cimicoidea: Vetanthocoridae) from the late Jurassic (Jung and Lee, 2012; Yao et al. 2006). In this analysis, we wanted to test if our molecular dating of the family Cimicidae is in concordance with oldest known cimicid fossil, *Quasicimex eilapinastes* Engel, 2008 from mid Cretaceous (ca. 100 MYA) (Engel, 2008). Our estimates placed the origin of Cimicidae at 93.8 MYA with a 95% highest probability density interval of 56–137 MYA (tree not shown). Accepting the fossil as a proxy for the minimum age of the Cimicidae, this clock estimate appeared as a reasonable result. To better account for variable evolutionary rates over the whole tree, we used the minimum age of *Q. eilapinastes* as an additional calibration point, setting a lognormal prior with a mean of 102.5 MY and standard deviation 0.06 MY for the diversification of the Cimicidae. The root in both analyses was given a weak uniform prior ranging from 0 to 350 MY.

We ran two successive MCMC chains with 100 million generations, sampling every 1000 generations. All chains had reached equilibrium at two million generations. When discarding 20% of the initial tree samples the consensus trees from each run produced the same topologies and the same branch support. We pooled samples from the two runs with the program “logcombiner” (Drummond et al., 2012) by discarding 50% of the initial trees from each run and computed a consensus chronogram based on 10000 resampled trees. Parameter estimates, including posterior probabilities and mean node ages with highest probability density intervals, were calculated in FigTree (Rambaut, 2014).

### Ancestral host character state reconstruction

We mapped ancestral host characters on the tree with time estimated nodes. We used Mesquite version 3.2 to prune the outgroup taxa from the tree and to collapse zero-length terminal branches. We coded terminal taxa with discrete trait characters according to the known host groups of each species: bats, birds (divided into Neoaves and Galloanseres) and humans. We then used the ‘trace ancestral character’ function to estimate ancestral states of nodes with maximum likelihood (Fig. 3A). A simple one-parameter Markov model (Drummond et al., 2012) was applied with these calculations, estimating the rate of state changes directly from the data (Maddison and Maddison, 2017). In a second approach, we coded terminal taxa with the discrete trait characters ‘specialist’ or ‘generalist’ (Fig. 3B). We then used the ‘trace ancestral character’ function to estimate ancestral states of nodes with maximum parsimony.

In addition, we inferred ancestral states at ancestral nodes using the full hierarchical Bayesian approach (integrating uncertainty concerning topology and other model parameters) as described in Huelsenbeck and Bollback (2001)) and integrated in MrBayes 3.2. The ancestral host character for the selected lineages (i.e. Cimicinae and (Haematosiphoninae + Cacodminae) at the KT boundary (the time of their assumed first colonization of bats) was also inferred using the full hierarchical Bayesian approach in MrBayes 3.2. All terminal taxa not belonging to one of these two lineages were coded as character “unknown host”.

### Ancestral spermalege character state reconstruction

Host characters for ancestral state reconstruction were mapped onto the dated tree. We used Mesquite version 3.2 to prune the outgroup taxa from the tree and to collapse zero-length terminal branches. We coded terminal taxa with discrete trait characters according to the position of the spermalege: 1) anterior-posterior body axis, i.e. segmental position (separate for tergites and sternites), and, 2) left-right body axis, i.e. a more left, middle (center) or right position. We then used the ‘trace ancestral character’ function to estimate ancestral states of nodes with maximum likelihood. A simple one-parameter Markov model (Lewis, 2001) was applied with these calculations.

## Supplementary information

Supplementary Information 1: Table with list of samples of the 34 species, covering 30% of extant species described to date from 6 out of 7 recognized subfamilies, or 17 out of 26 genera described to date (Henry, 2009), as well as their collectors and the GenBank accession number.

Supplementary Information 2: Graph showing host usage of the Cimicdae, reconstructed using a strict definition of generalism.

Supplementary Information 3: Evolutionary occurrence of extant bedbug lineages and their host genera, as extracted from our phylogenetic tree.

Supplementary Information 4: Host relationships (tanglegram) of Cimicidae parasitic on bats.

Supplementary Information 5: Figure: Host relationships (tanglegram) of the bird-parasitic Haematosophinae.

Supplementary Information 6: Supplementary discussion. Three issues are briefly discussed: Heterotopy of reproductive structures in the animal kingdom, evolution of mating position in the Cimicidae and paraphyly of *Cimex* and former *Oeciacus*.

Supplementary Information 7: Cross-species comparison of morphological variation in the copulatory organs of bedbugs.

Supplementary Information 8: Data file: Summary of nodes and character changes in the Cimicidae

Supplementary Information 9: Characteristics of molecular markers used for the analyses.

Supplementary Information 10: Bayesian analysis (BA) of phylogenetic relationships of the Cimicidae inferred from individual genes. Consensus trees inferred from the single gene fragments (18S rDNA part1 and part 2, COI, 16S rDNA, 28S D3 rDNA

Supplementary Information 11: MrBayes consensus tree using one representative species of the closest phylogenetic taxa (e.g. Anthocoridae, Nabidae and Plokiophillidae) within our outgroup sampling.

Supplementary Information 12: List of primers used and PCR conditions used in the study.

Supplementary Information 13: Maximum Likelihood analysis of the combined molecular data set.

Supplementary Information 14: GBlock alignment tests for trees using strict and relaxed models.

Supplementary Information 15: Full alignment file.

Supplementary References

## Statement on authorship

SR, MS-J, EHM and KR conceived the study. SR, OB, ODI, MS-J, PB, OC, EF, MM, RN, NS, EHM, FAAK, MPL and KR did fieldwork and extensively contributed material or sequences. SR carried out the molecular work. SR, EW and KR analyzed the data. SR and KR wrote the first draft, SR, OB, MS-J, MLP, EHM, EW, and KR carried out the first revision. All authors, except ODI contributed to all subsequent revisions.

## Acknowledgments

We thank E. Vargo, M. Stoneking, M. Lehnert and N. Tatarnic for comments on the manuscript, all people mentioned in table S1 for providing samples and L. Lindblom, K. Meland, D. Rees for help with molecular work, and R. Mally and B. Jordal for help with data analysis. For specimen loan or opportunity for inspection or DNA extraction we thank Museum Senckenberg Frankfurt, Naturalis Biodiversity Center Leiden, Natural History Museum London, Tel Aviv University Department of Zoology, Universidad Nacional Autónoma de México Instituto de Biología, Universiti Malaysia Sarawak Zoology Department, Field Museum Chicago and University of Texas Insect Collection. The Forest Department of Sarawak provided permits (Research: #NCCD 907.4.4 (JLD.12)–85; Park # 209/2015 and Export #16001).

## Additional information

**Funding**

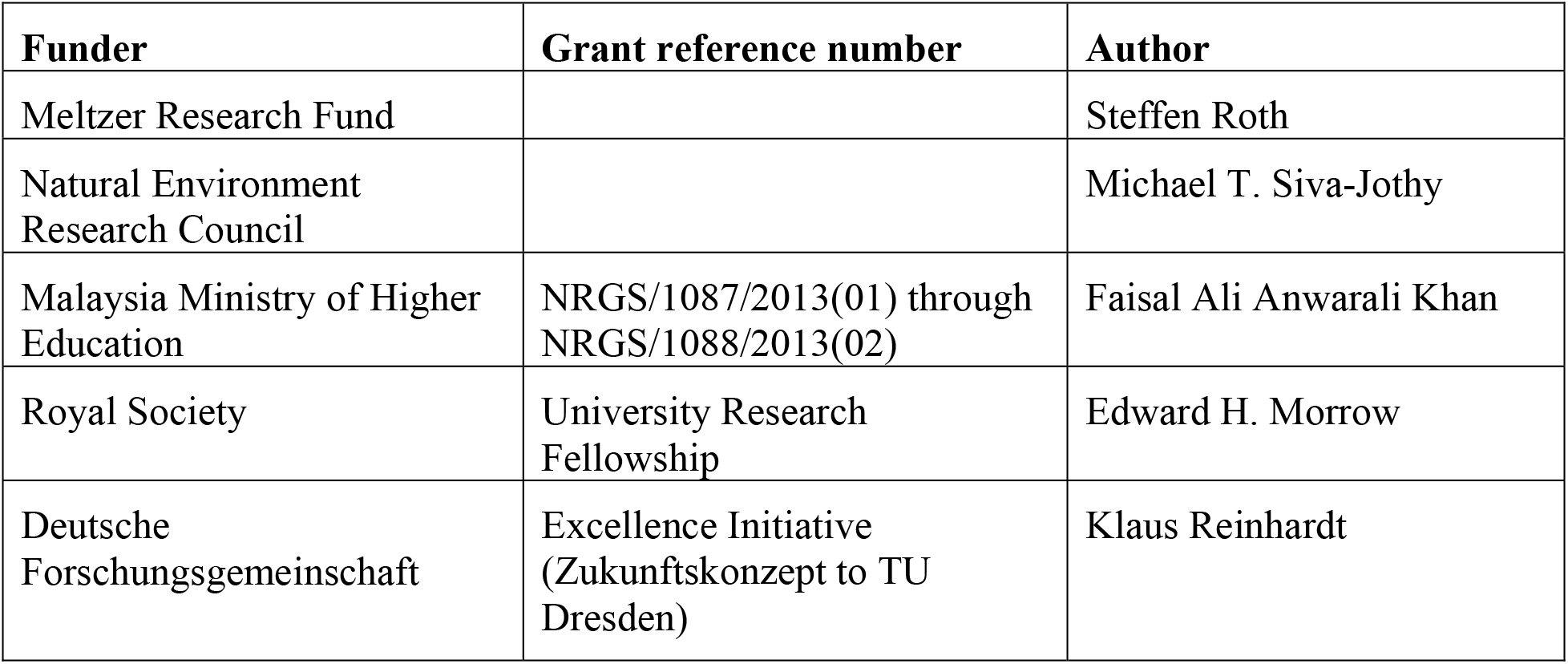

**Author contributions**

**Author ORCIDs**

**Data availability**

Source files for the phylogenetic analyses of Figures 1 and 2 and Supplementary Information 10, 11 and 14 have been uploaded. They will be made available via dryad.

## Supplementary Information 1 to 15

**Supplementary Information 1:**

List of samples of the 34 species, covering 30% of extant species described to date from 6 out of 7 recognized subfamilies, or 17 out of 26 genera described to date (Henry, 2009).

*- See separate file -*

**Supplementary Information 2:**
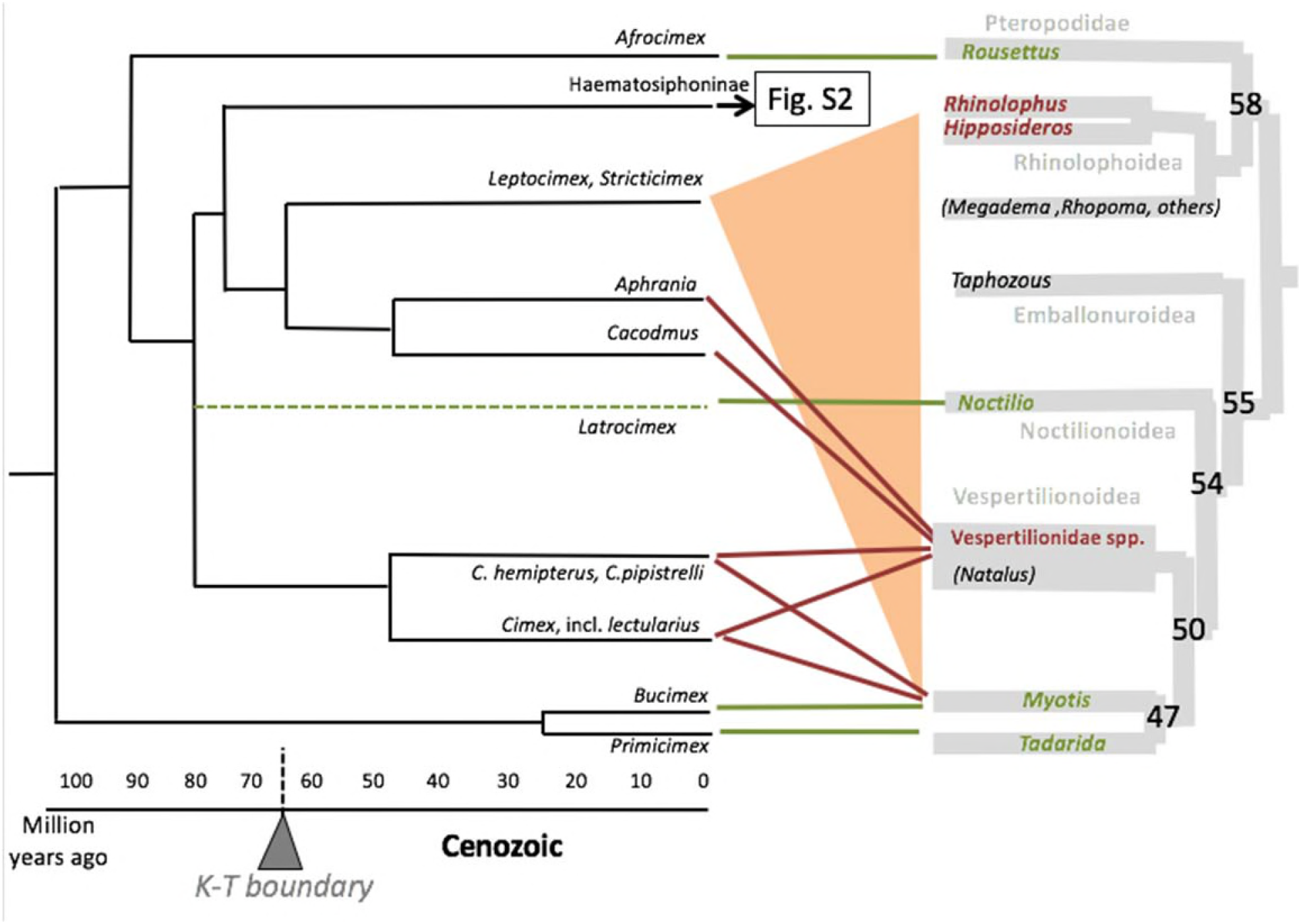
Host reconstruction using a stricter definition of generalism. Here, host generalism is defined as utilizing more than three host genera. The host spectrum was obtained from the same sources as for figure 3, with an additional record for C. *sparsilis* on domestic dog (Coetzee & Segerman, 1992).

**Supplementary Information 3:**
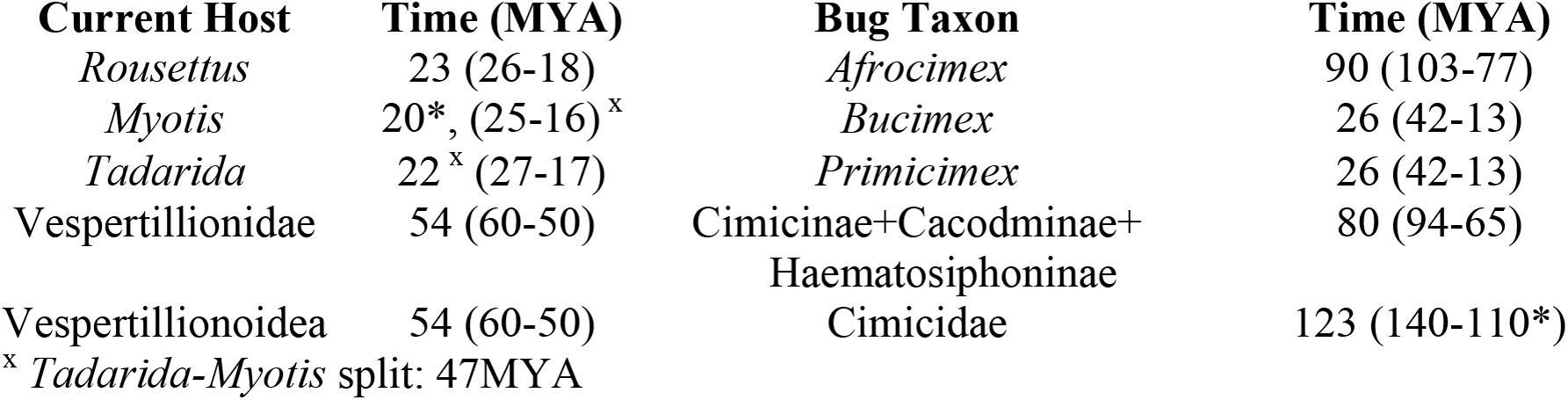
Evolutionary occurrence of extant bedbug lineages and their host genera, as extracted from our phylogenetic tree. (*) indicates molecular ages which are confirmed by oldest fossils (less than ±10 MYA). Mean age, 95% lower and upper highest posterior distribution inferred by BEAST (Drummond et al., 2012) is reported.

Event-, distance- or topology-based cophylogenetic tests were not applied because the molecular and phylogenetic resolution of host and parasite trees did not match and because the few exhaustive host species lists of the Cimicidae that exist (Usinger, 1966, Ueshima, 1968; Di Iorio et al., 2013); Országh et al., 1990; Coetzee & Segerman, 1992; Supplementary Information 1) suggest that over-precision should be avoided.

**Supplementary Information 4:**
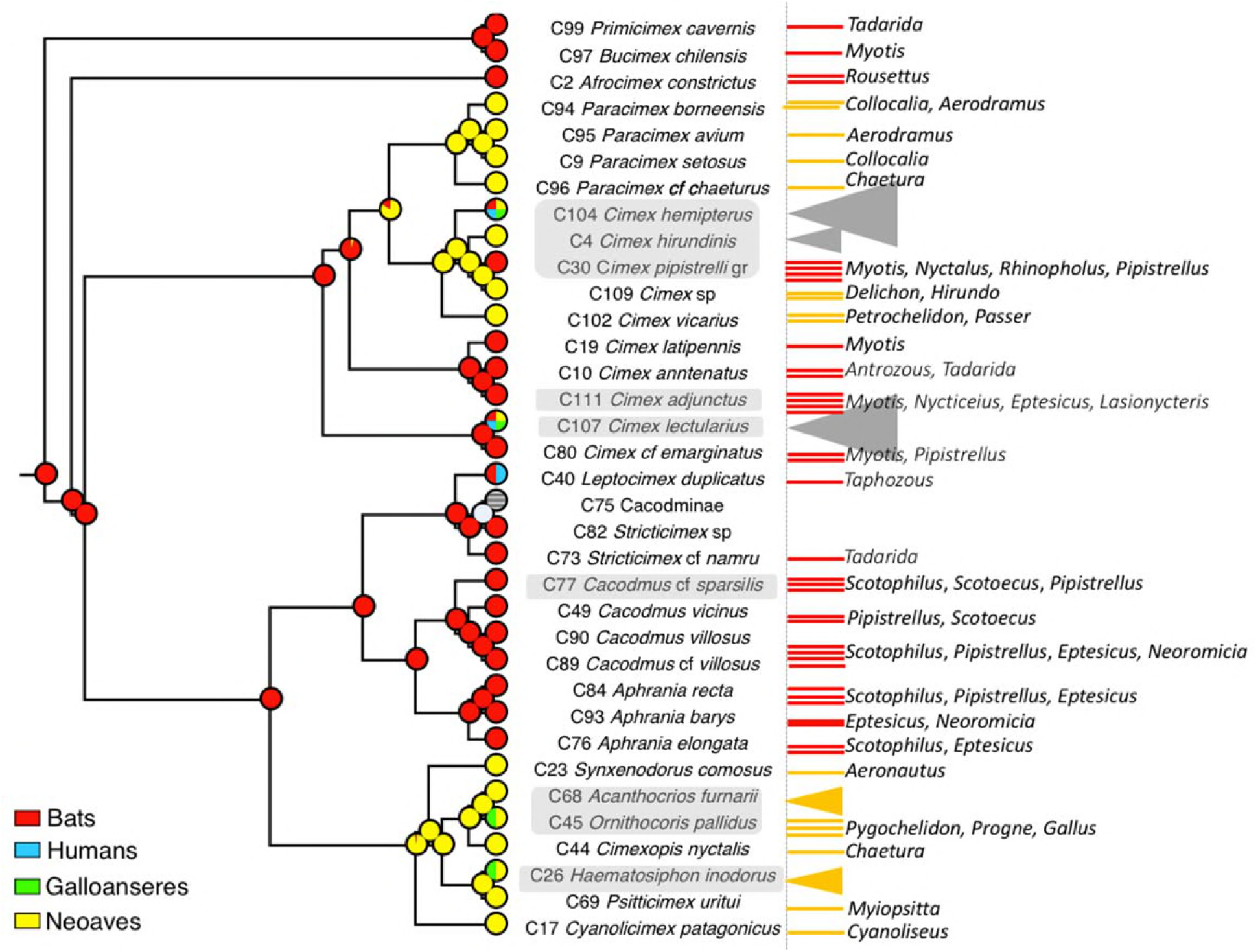
Host relationships (tanglegram) of Cimicidae parasitic on bats. Specialists having only one species or genus as hosts are shown with green connectors, generalists with a wider range of host taxa are shown with red connectors; *Leptocimex* and *Stricticimex* utilize hosts except *Noctilio* that phylogenetically are wide apart (orange). Bat phylogeny according to Teeling (2005), host spectrum after (Usinger, 1966; Ueshima, 1968; Di Iorio et al., 2013); Országh et al., 1990; Supplementary Information 1).

**Supplementary Information 5:**
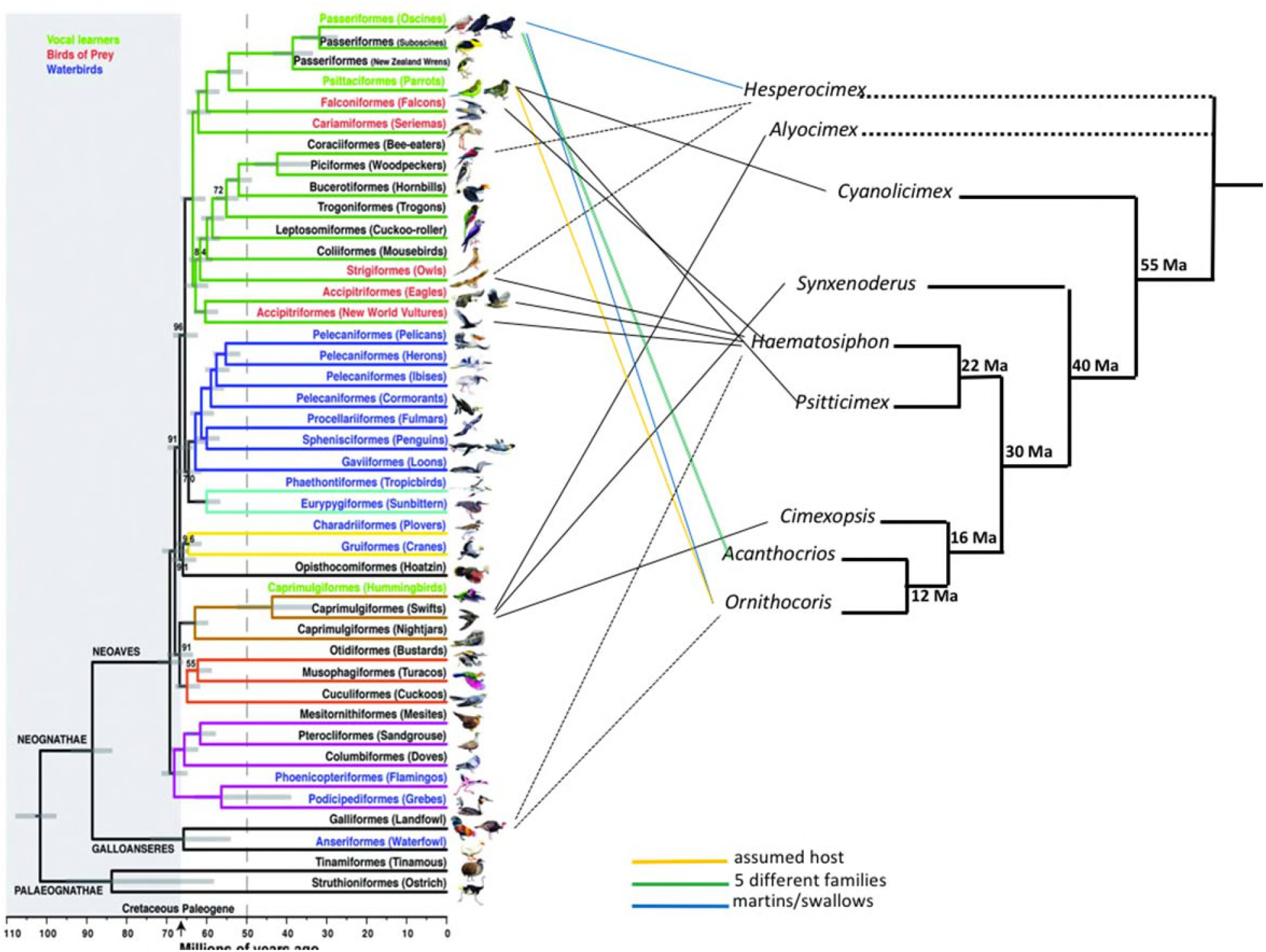
Host relationships (tanglegram) of the bird-parasitic Haematosophinae. Primary hosts (solid line) and secondary hosts (long dashed line) (after Di Iorio et al., 2010). Dotted branches are species that were not analysed in our study. The Haematosiphoninae (diverged around 50 MYA) and the bird-parasitic *Paracimex* (around 15 MYA) or *Cimex vicarius* (around 18 MYA) all appeared long after their respective swift or swallow host groups had appeared in the early Eocene (Brown et al., 2008; Ericson et al., 2014). Phylogram of birds from Jarvis et al. (2014). Hosts were compiled from (Usinger, 1966, Ueshima, 1968; Di Iorio et al., 2013; Országh et al., 1990; Supplementary Information 1).

**Supplementary Information 6**

**Discussion.**

**Heterotopy of reproductive structures in the animal kingdom**. The two other examples where substantial heterotopy of reproductive structures is found concerns the site of traumatic intromission in *Siphopteron* sea slugs (Lange et al., 2014) and the genus-specific position of the genital opening in female mites (Lee, 1970).

**Mating position in the Cimicidae**. Cimicidae show two novel traits derived from the ancestral neopteran female-above mating position: the false male-above position in the front inseminators and the male-above position in the back inseminators (Usinger, 1966). Unfortunately, the variation in male mating position is not documented for individual bedbug species.

A female morphological response to changes in male mating behavior would be consistent with female genitalia duplication which occurred twice independently in the Cimicomorpha, in *Cardiastethus limbatellus* (Anthocoridae), and the Plokiophilidae (Schuh & Slater, 1995; Carayon, 1977), as well as with the existence of a spermalege in males of a species with male-male inseminations (Reinhardt et al., 2007).

**Paraphyly of *Cimex* and formerly *Oeciacus***. Relating fittingly to our dedication, Usinger (Usinger, 1966) had already shown substantial reproductive isolation between the European and North American martin bug species *C. hirundinis*, and *C. vicarius* (previously *Oeciacus*), compared to other cimicid species. Upon these results, he developed a novel, now rather modern, species concept of stepwise postmating isolation. Ironically, had he followed his own concept more than traditional morphological similarity, which is now known to be partially host-dependent (Balvin et al., 2013), he would have reached the conclusion of a paraphyly of the former genus *Oeciacus* that the current molecular analysis arrived at.

**Supplementary Information 7:**
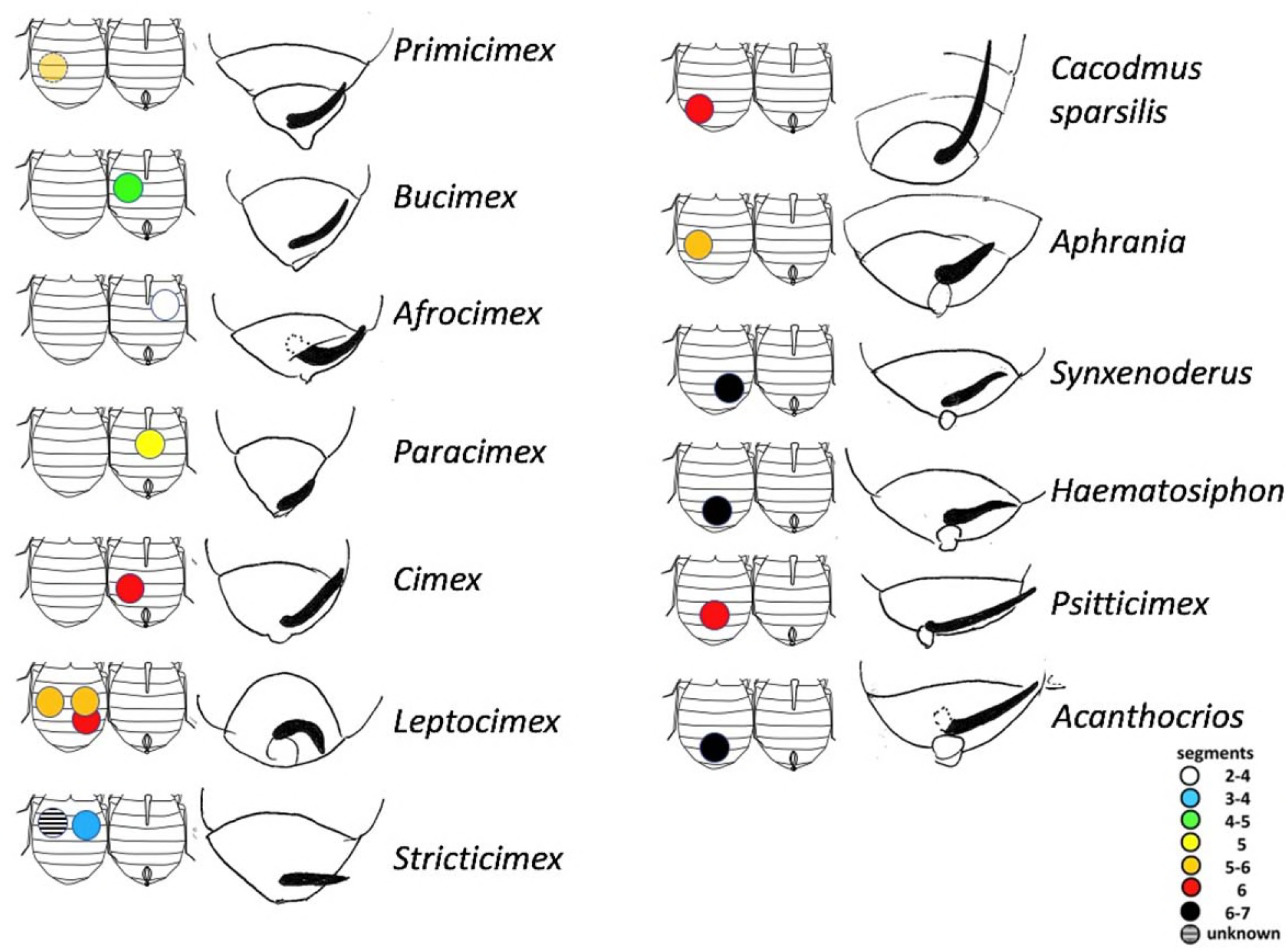
Cross-species comparison of morphological variation in the copulatory organs of bedbugs. The genus or species name is given in the right column, the female abdomen on the left column (dorsal side left, ventral side attached to the right). The segment number is color-coded as in figure 2, and the position indicated by the coloured dots. The intromittent organs of male bedbugs are shown in black (redrawn from Usinger, 1966) are shown on the right of each panel. The male organs are not to scale, only the last abdominal segments are depicted (for size comparison cf. the last abdominal segments of in female drawings).

**Supplementary Information 8: Summary of nodes and character changes in the Cimicidae** *See separate file*

**Supplementary Information 9:** Characteristics of molecular markers used. To implement Kimura’s two-parameter model (K2) in BEAST 1.8.4, we selected the Hasegawa-Kishino-Yano (HKY) model and set “base frequencies” to “All Equal”. For many taxa sampled, the two 18S fragments did not overlap. Therefore, the two fragments were analyzed separately.

| Gene | Sequence length (bp) | Number of missing taxa | Alignment position | Parsimony informative | Variable sites | Evolution model |
| --- | --- | --- | --- | --- | --- | --- |
| COI | 591-659 | 2 | 659 | 335 | 359 | GTR+G+I |
| 16S rDNA | 361-519 | 1 | 571 | 311 | 395 | TN93+G+I |
| 28S rDNA | 301-337 | 1 | 363 | 82 | 121 | K2+H |
| 18S rDNA part 1 | 561-988 | 2 | 1121 | 266 | 415 | K2+G+I |
| 18S rDNA part 2 | 598-697 | 3 | 711 | 78 | 144 | K2+G+I |

**S10a.**
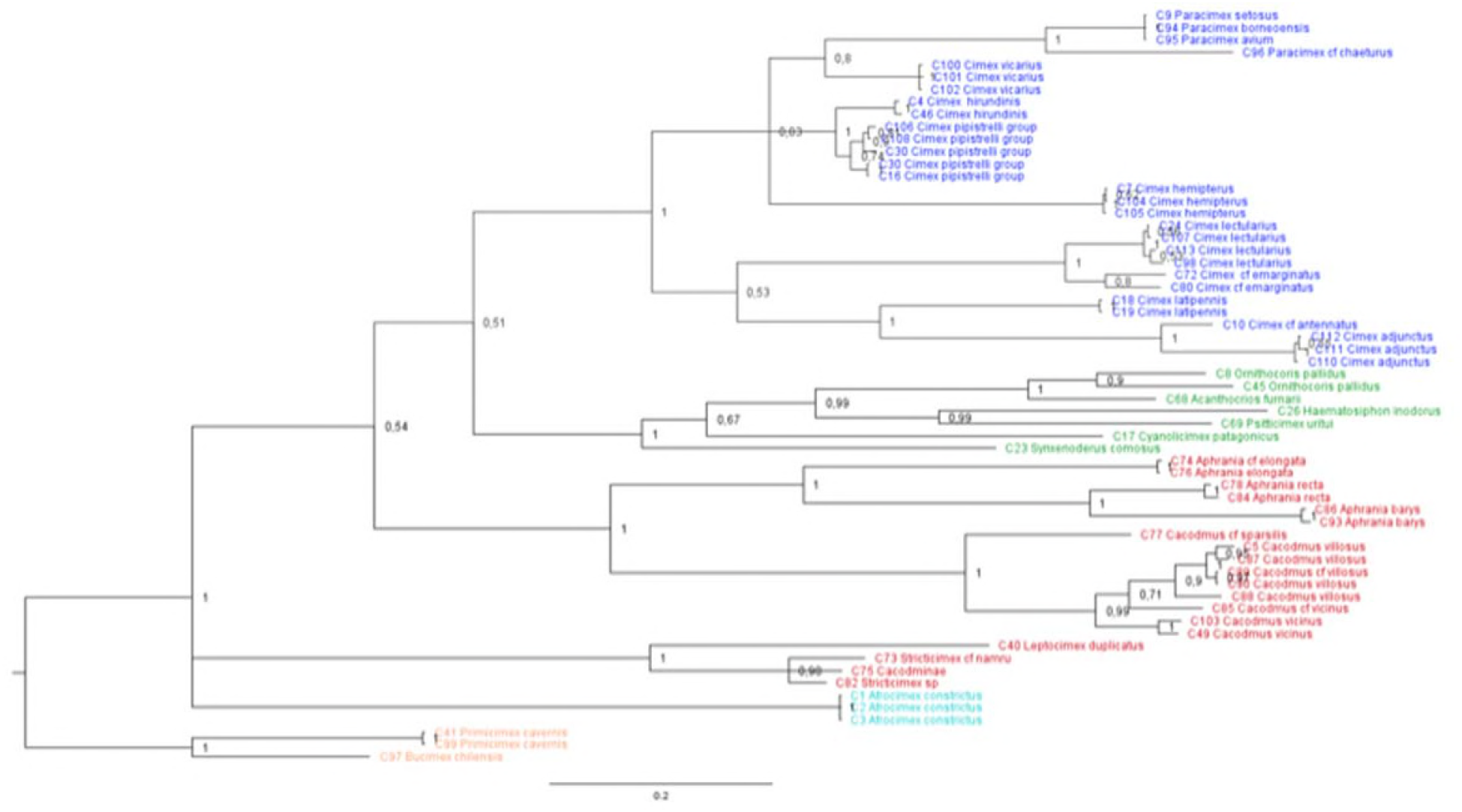
Bayesian Analysis of the COI gene. The underlying data (***Additional Data 2***) are available at ***dryad*** (see additional files; currently uploaded for review).

**S10b.**
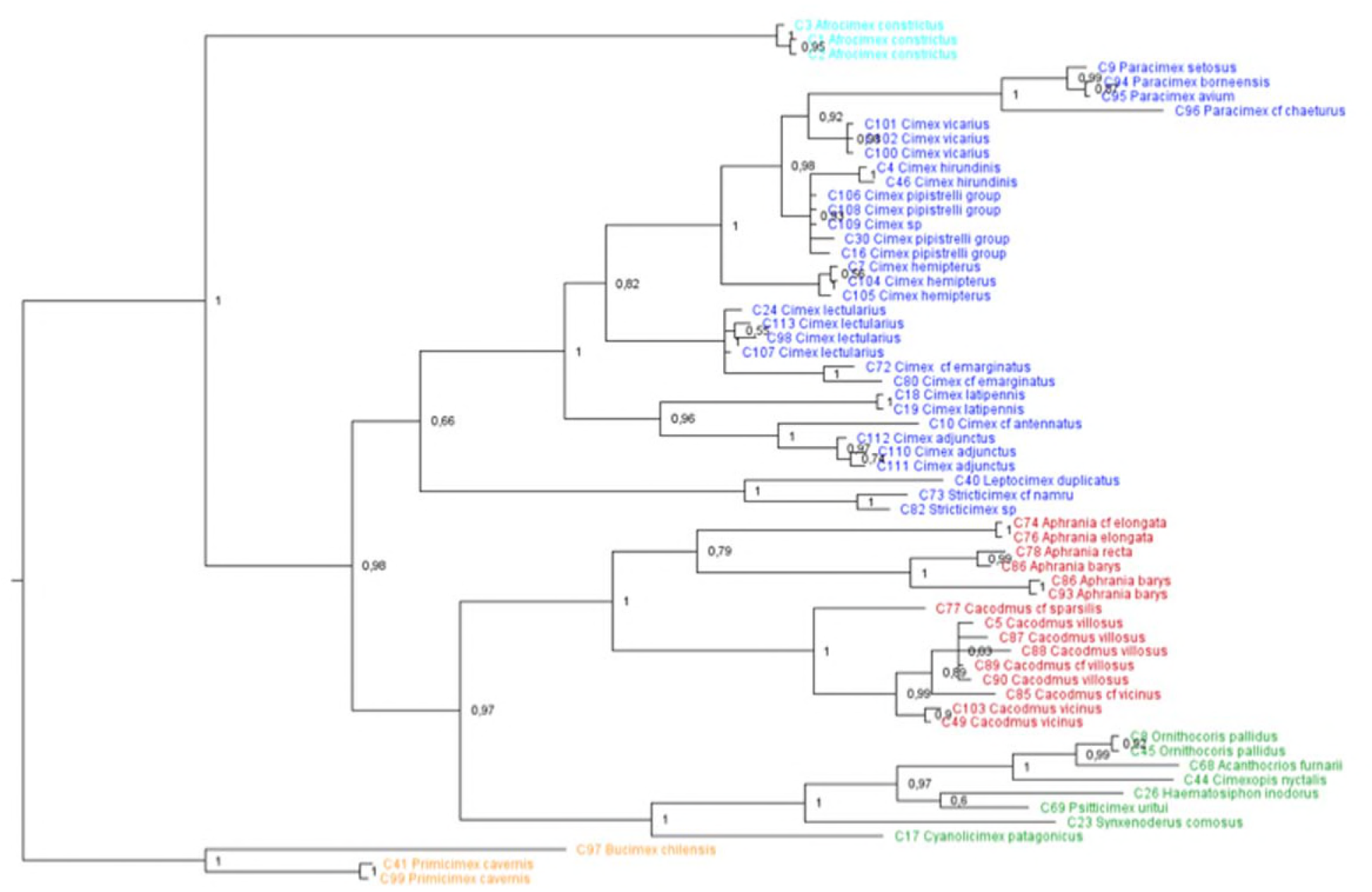
Bayesian Analysis of the 16S rDNA gene. The underlying data (***Additional Data 3***) are available at ***dryad*** (see additional files; currently uploaded for review).

**S10c.**
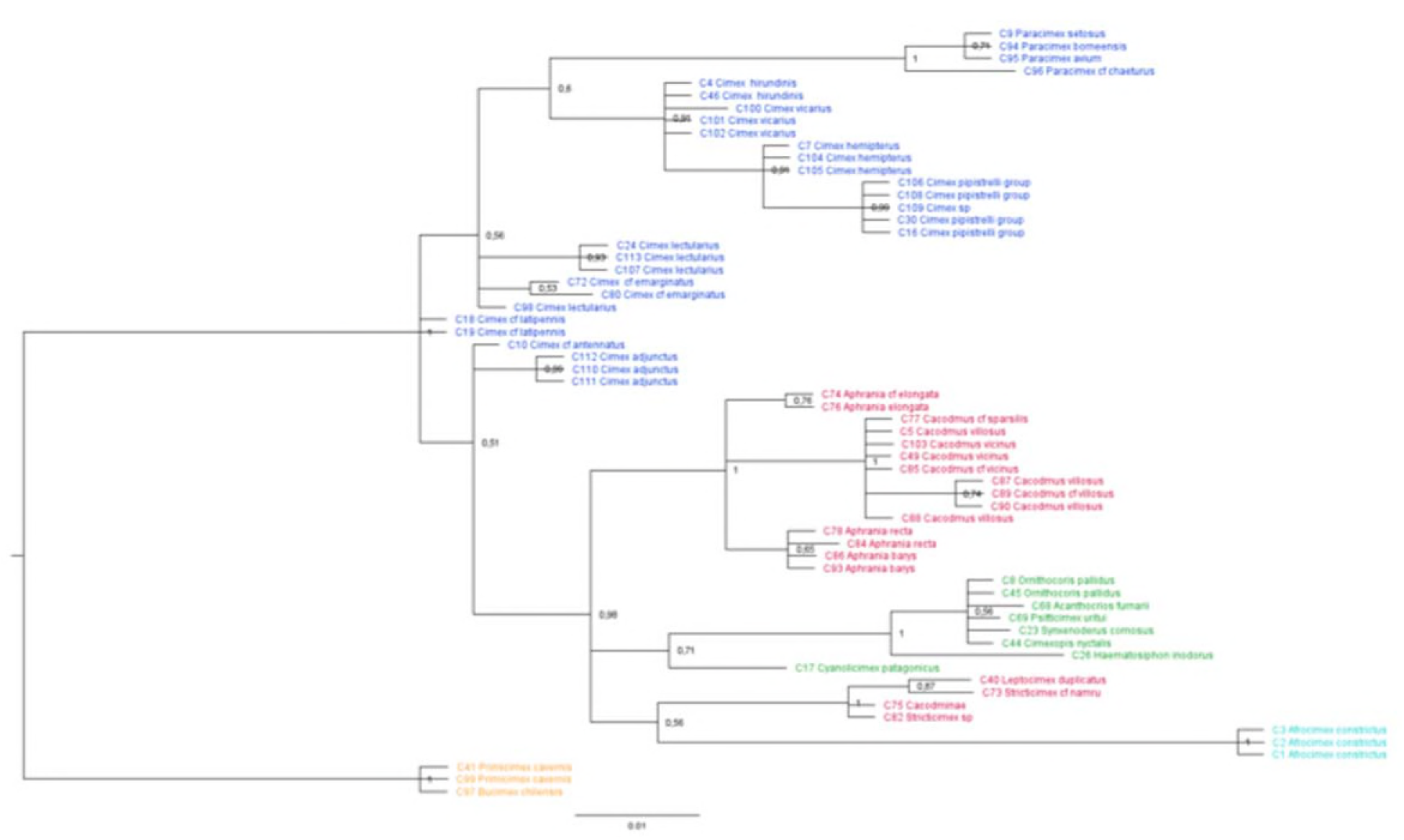
Bayesian Analysis of the 28S D3 rDNA gene. The underlying data (***Additional Data 4***) are available at ***dryad*** (see additional files; currently uploaded for review).

**S10d.**
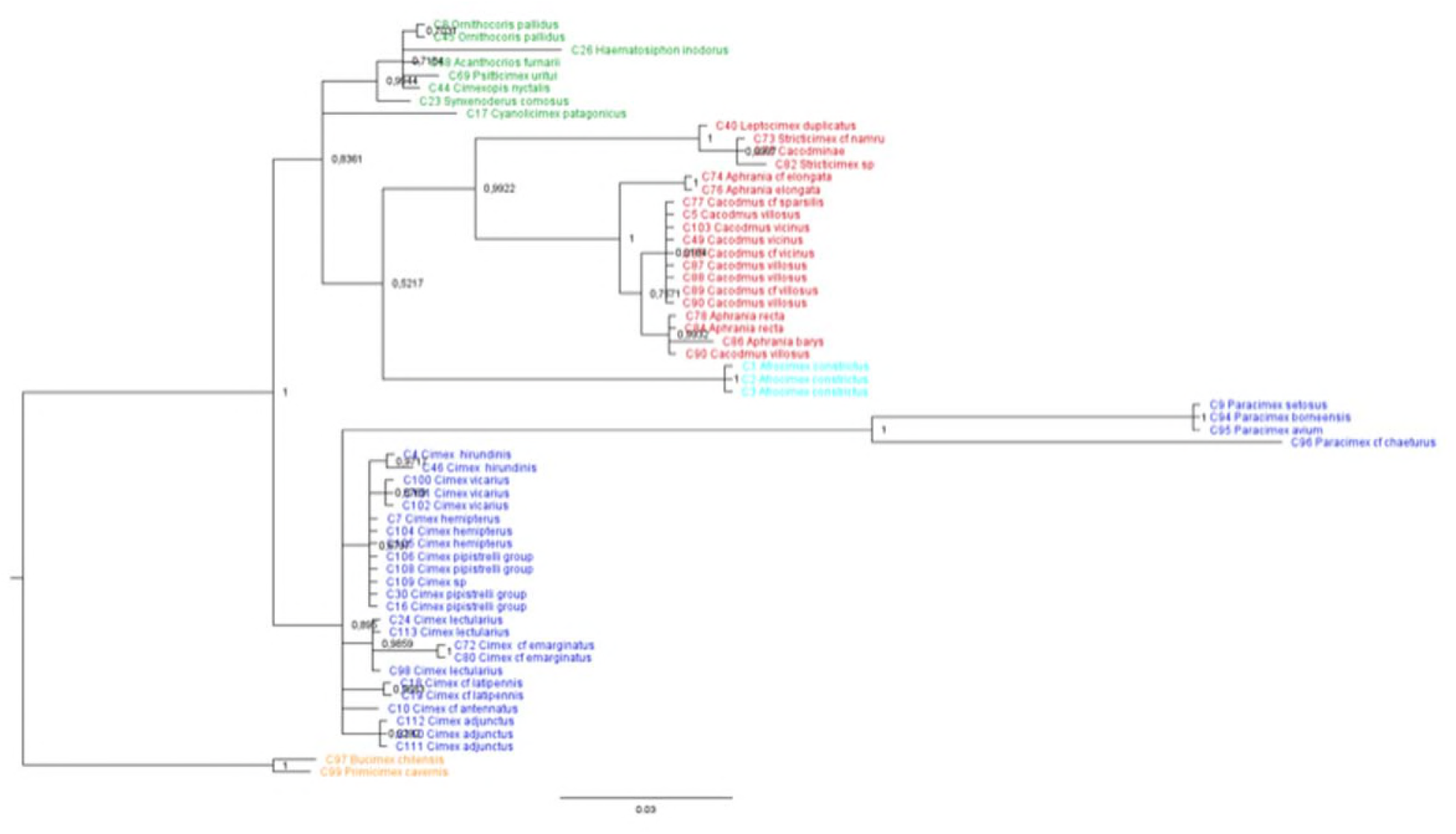
Bayesian Analysis of the 18S rDNA gene, part 1. The underlying data (***Additional Data 5***) are available at ***dryad*** (see additional files; currently uploaded for review).

**S10e.**
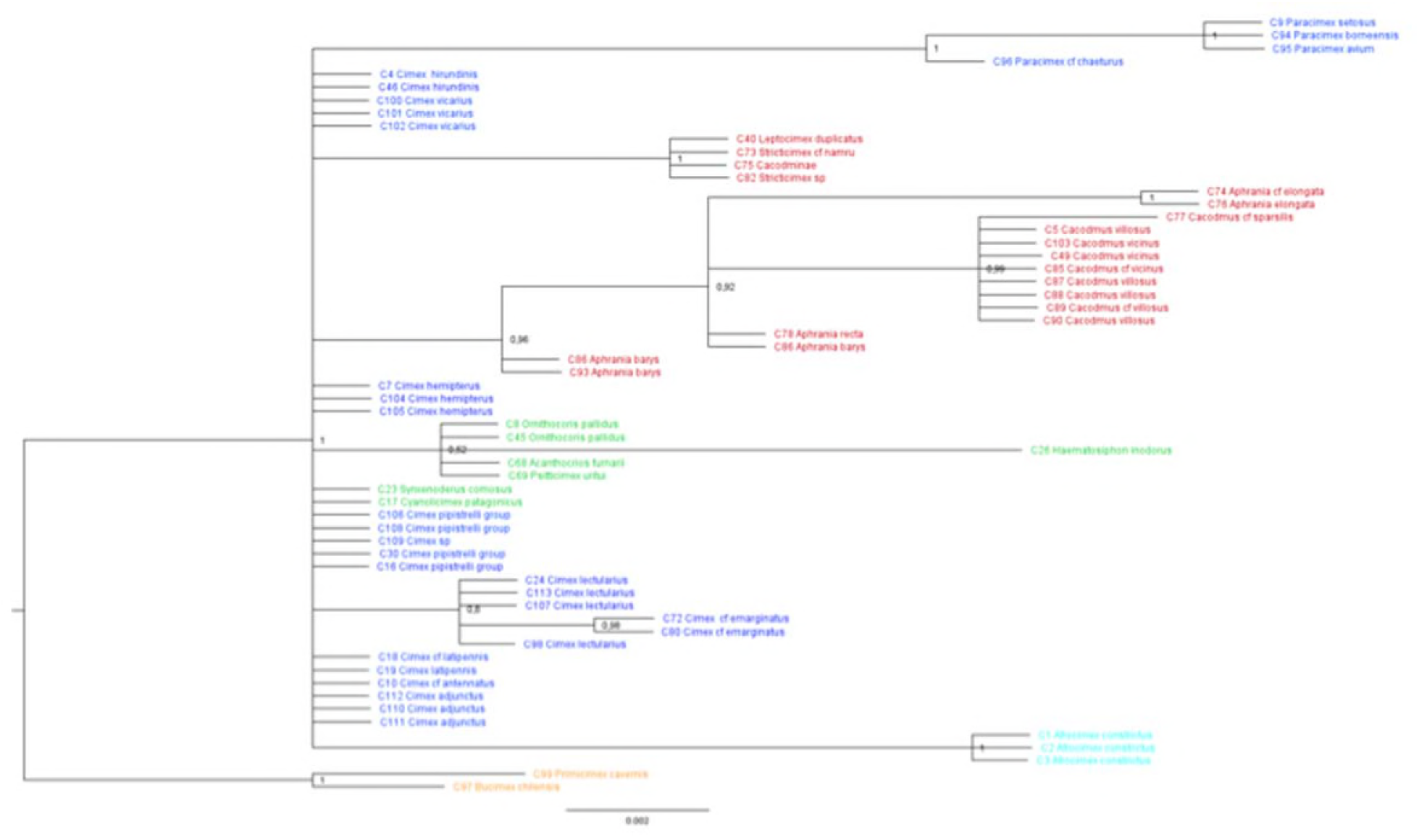
Bayesian Analysis of the 18S rDNA gene, part 2. The underlying data (***Additional Data 6***) are available at ***dryad*** (see additional files; currently uploaded for review).

**Supplementary Information 10:**

**Fig. a-e Bayesian analysis (BA) of phylogenetic relationships of the Cimicidae inferred from individual genes.** The analysis was carried out using MrBayes v.3.2.1 (Ronquist et al., 2012) for individual genes, substitution models were as chosen in the combined data set analysis (Supplement 13). Details for the settings in MrBayes for single genes will be available via dryad (see ***Additional Data 2–6***). Consensus trees inferred from single gene fragments (COI, 16S rDNA, 28S D3 rDNA, 18S rDNA part1 and part 2) show their different phylogenetic information but also that single gene analyses are unable to recover phylogenetic relationship. **a**) COI, **b**) 16S rDNA, **c**) 28S D3 rDNA, **d**) 18S rDNA, part 1, **e**) 18S rDNA, part 2.

**Supplementary Information 11:**
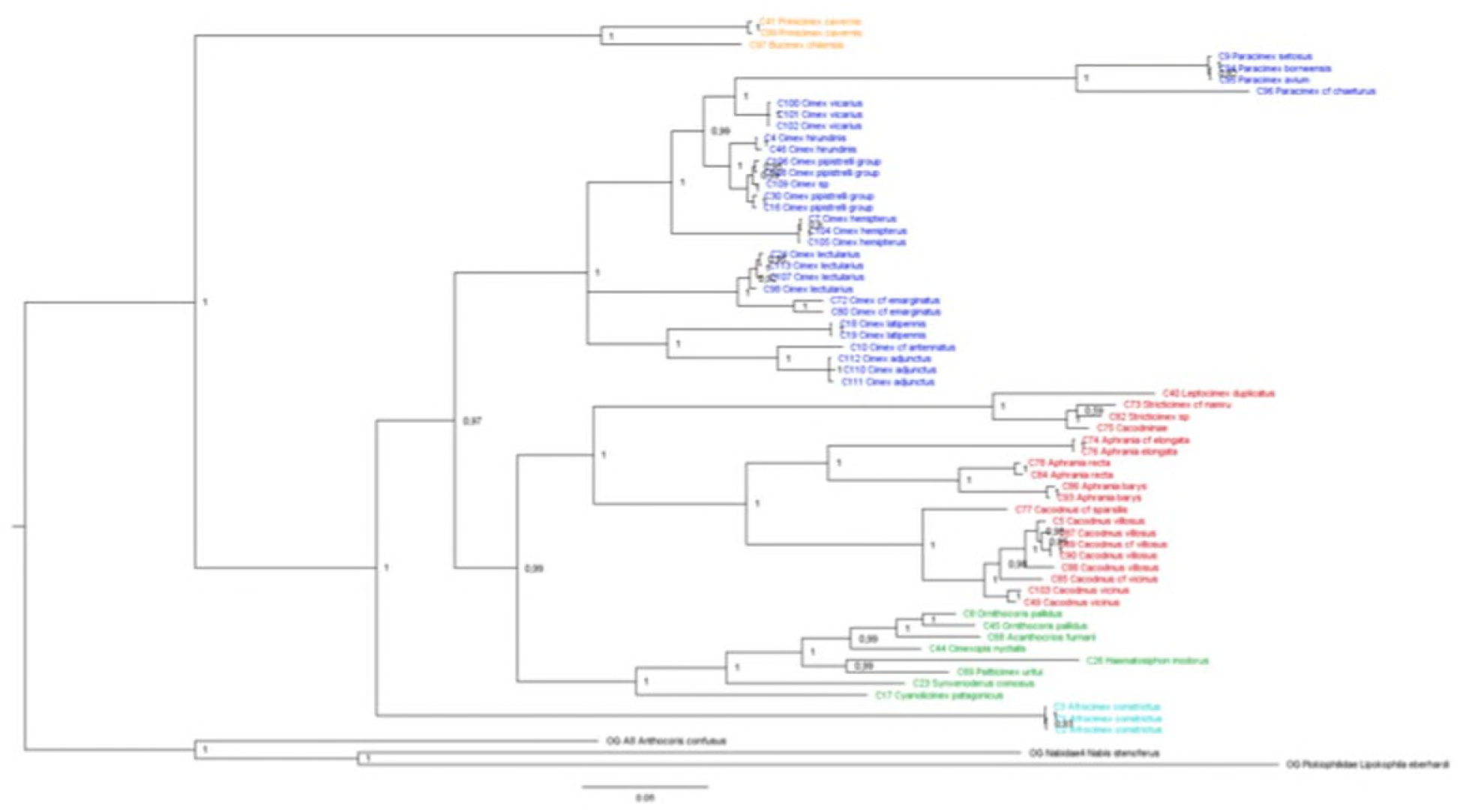
MrBayes consensus tree using one representative species of the closest phylogenetic taxa (e.g. Anthocoridae, Nabidae and Plokiophillidae) within our outgroup sampling. The tree is a Bayesian consensus tree based on four genes (see Material & Methods). Numbers beside the nodes indicate posterior probability values. Topology and support value of the Cimicidae clades did not change due to different outgroup sampling (see Figure 1). The underlying data (***Additional Data 7***) are available at ***dryad*** (see additional files; currently uploaded for review).

**Supplementary Information 12:**
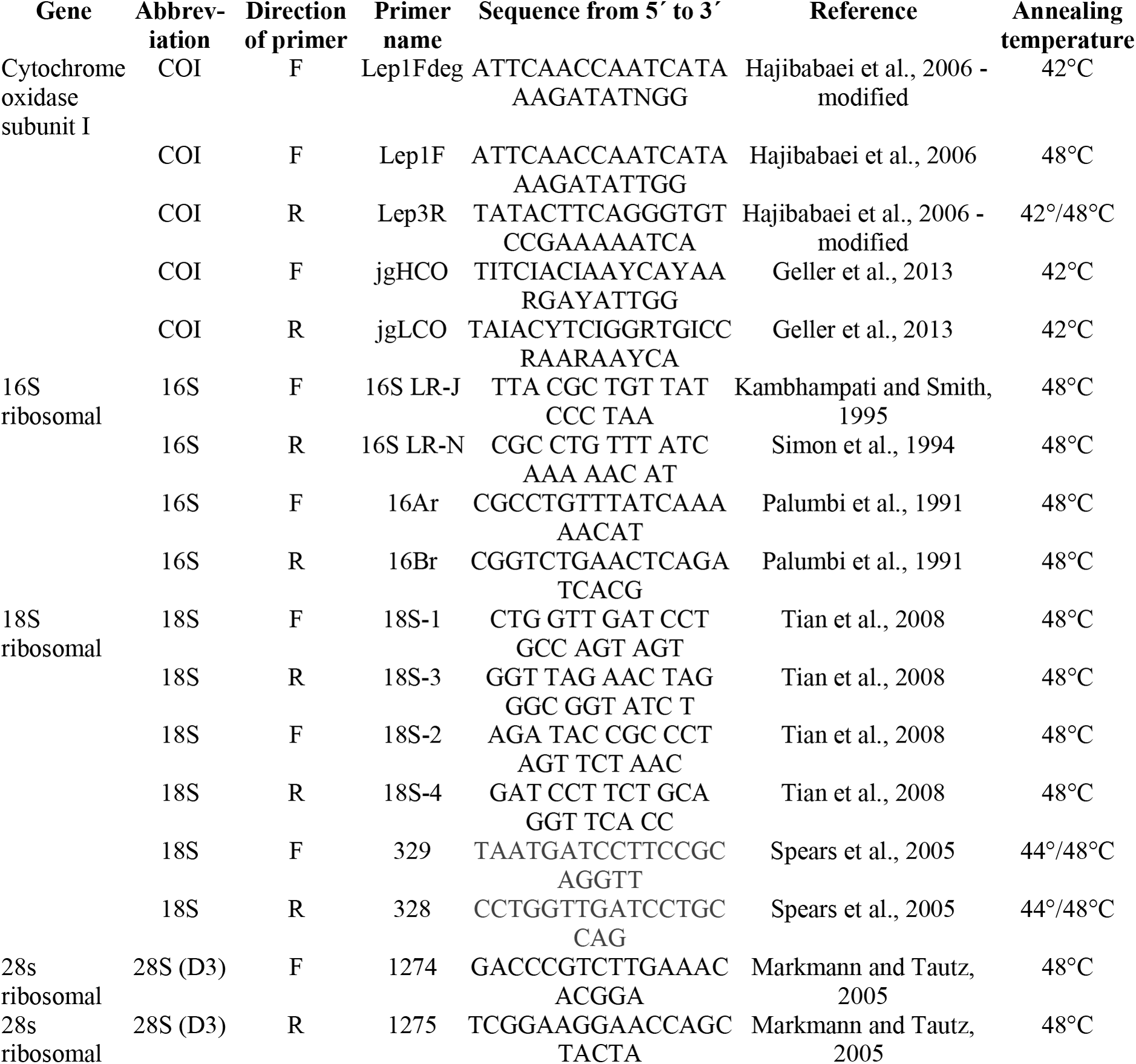
List of primers used and PCR conditions.

**Supplementary Information 13:**
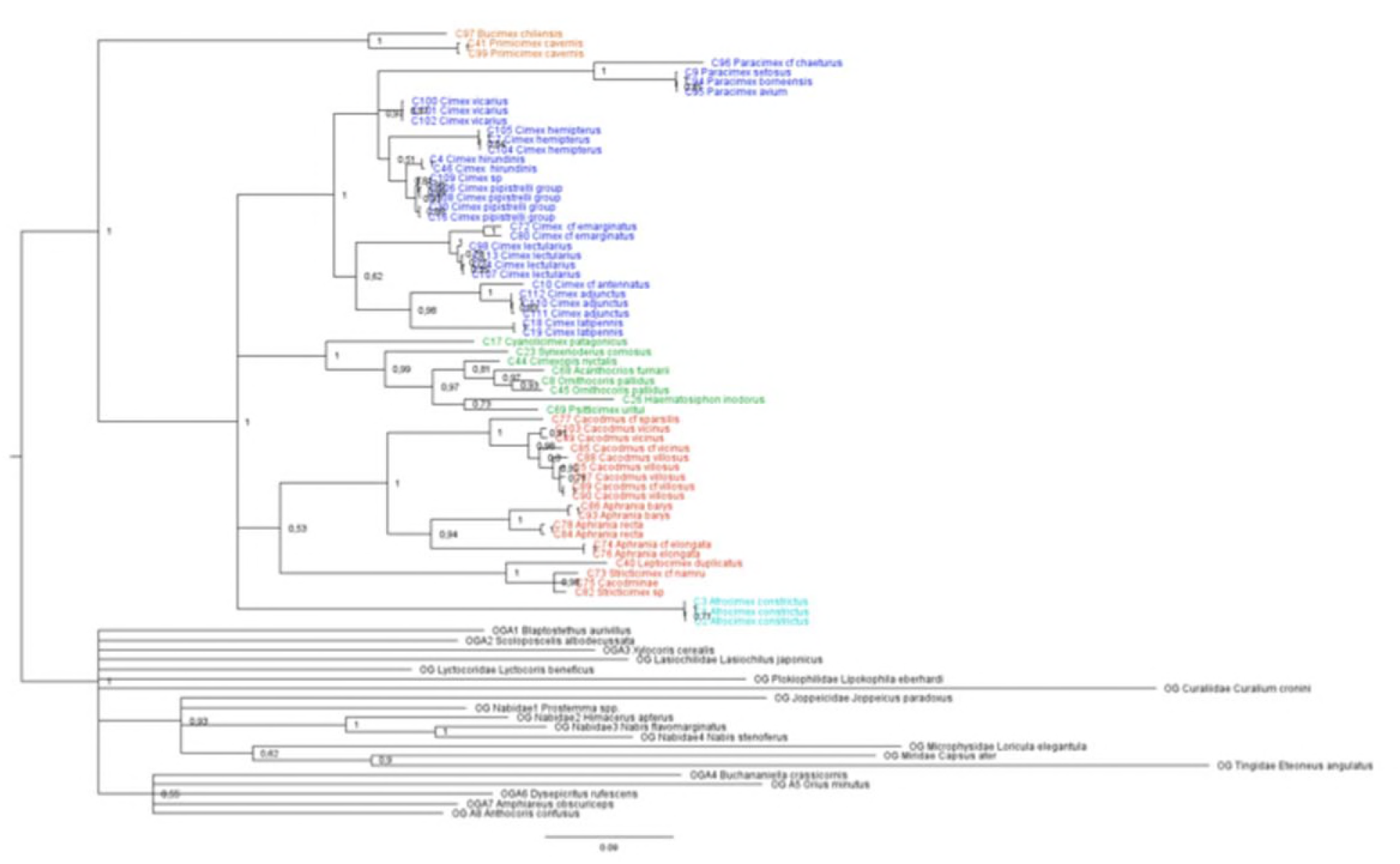
Maximum Likelihood analysis of the combined molecular data set. The Maximum Likelihood analysis confirmed the results of the BA (fig. S1) but the sister relationship of Cacodminae and Haematosiphoninae was not resolved. There was also low support for the node (*Leptocimex*+*Stricticimex*) + (*Aphrania*+*Cacodmus*). The underlying data (***Additional Data 8***) are available at ***dryad*** (see additional files; currently uploaded for review).

**Supplementary Information 14a:**
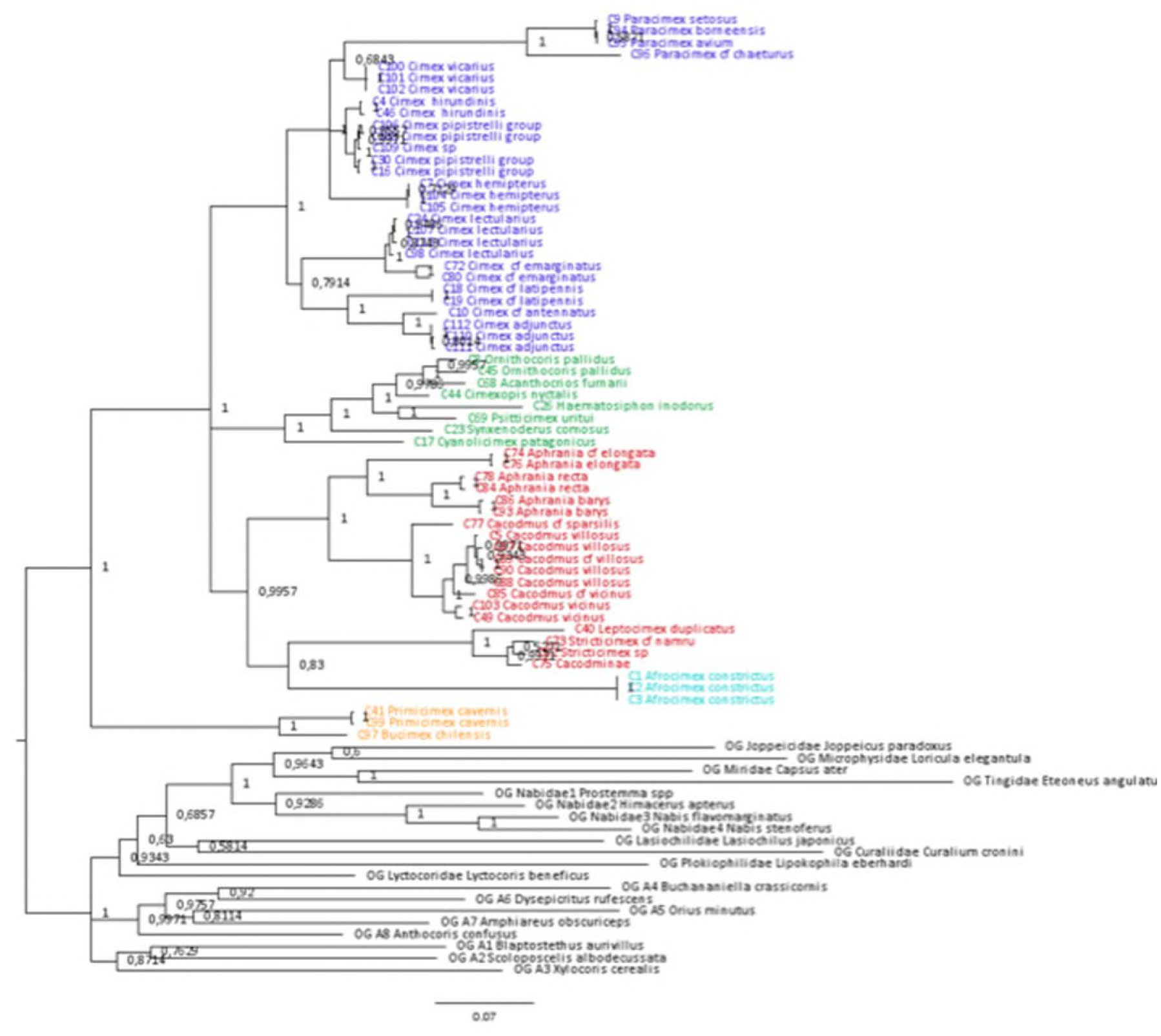

**Supplementary Information 14b:**
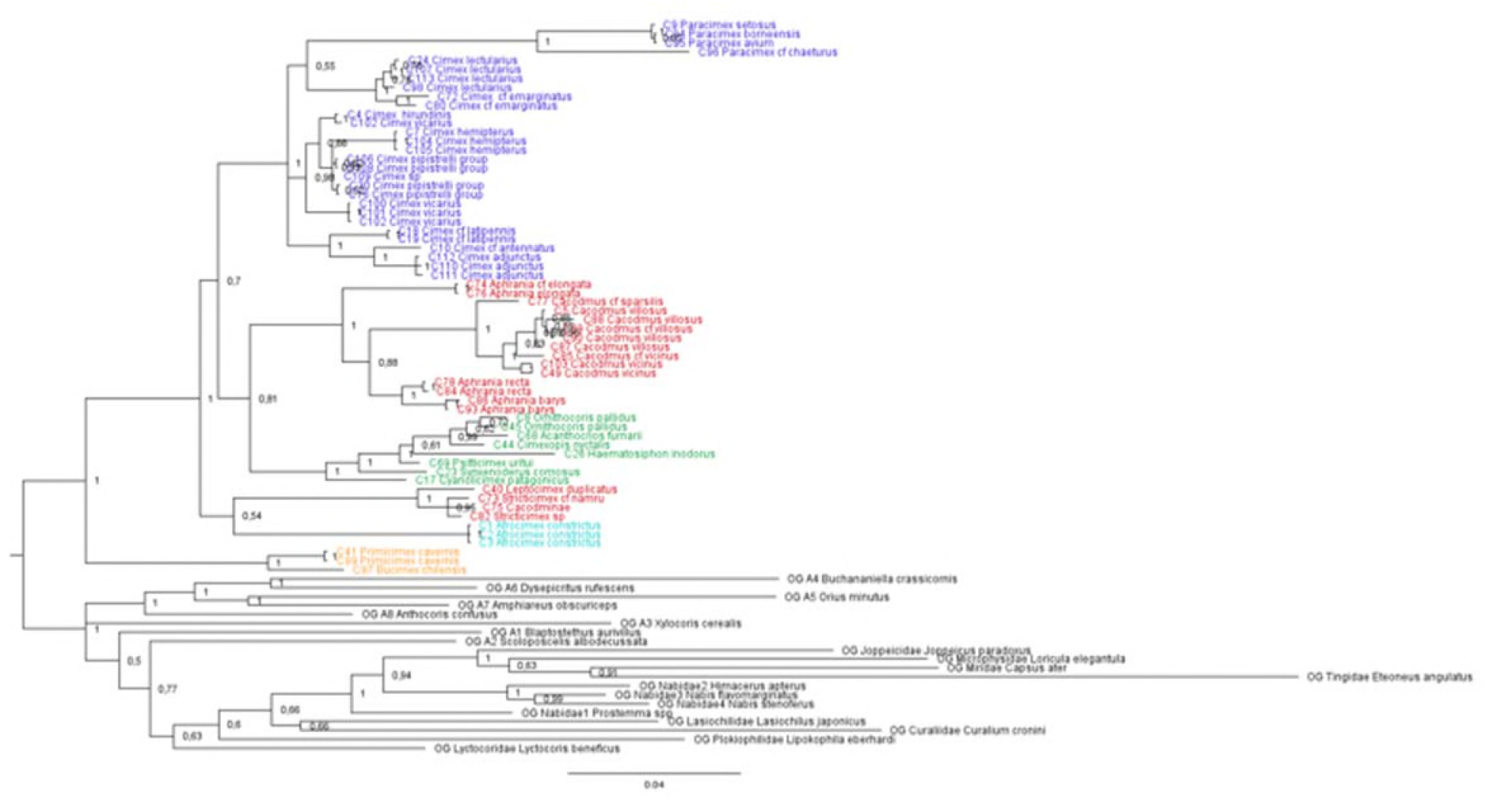

**Supplementary Information 14:**

**GBlock alignment tests for trees using strict and relaxed models.** Neighbor Joining (NJ) tree for the combined data set with original alignment set and GBlocks data set with tree strict (a) and relaxed (b) model using default settings of Gblocks V.0.91b (Castresana, 2000). NJ analysis was performed in MEGA v.6 (Tamura et al., 2013). NJ analysis using strict (a) and relaxed GBlock alignments (b) of all gene markers separately showed no significant effect of alignments and no need to eliminate poorly aligned positions and divergent regions, except some outgroup taxa. The original alignment data set was used for further analysis. Samples C41 and outgroup taxa *Curalium cronini* were removed from this analysis because of missing sequences.

The underlying data (***Additional Data 9 for 14a and 10 for 14b***) are available at ***dryad*** (see additional files; currently uploaded for review).

**Supplementary Information 15: Alignment file**.

*See separate file*

